# RhabdoForge: A Modular, Biophysically-Grounded Rendering Framework for Insect Vision Neuroethology

**DOI:** 10.64898/2026.08.25.747007

**Authors:** Florent Le Moël, Barbara Webb

## Abstract

Insects solve complex behavioural tasks with remarkable efficiency, using minimal neural hardware tuned to the specific requirements of their ecological niches. To truly understand or replicate these behaviours, it is insufficient to model the brain in isolation: one must account for the dynamic, closed-loop interactions between the environment, the physical organisation of the sensory periphery, and internal biophysical dynamics. To address these issues for visually controlled behaviours, we present RhabdoForge, a modular, hardware-agnostic and high-performance rendering framework specifically designed for insect neuroethology and neuromorphic research. Designed for seamless integration into Python-based workflows, RhabdoForge implements both real-time ray-tracing and stochastic path-tracing using hardware-agnostic GPU pipelines. Crucially, the engine moves beyond the static “ommatidium-as-a-pixel” paradigm by introducing a fully parametrisable model where every layer of the compound eye (from the geometric shape and the topological lattice to the internal rhabdomere blueprint) is a discrete, swappable component. The engine is capable of simulating the high-frequency, sub-ommatidial rhabdomere photomechanical actuation, waveguide optics, intracellular pupil adaptation, and retinal sheet movements, allowing for the investigation of a variety of active sensing phenomena within a real-time closed-loop environment. The framework also includes an automated morphological pipeline that allows transforming 2D anatomical data into faithful 3D sensory models. We demonstrate the engine through two case studies: a closed-loop optic-flow centring response in a virtual tunnel, and a decomposition of the mechanisms that can contribute to spatial hyperacuity (axial and lateral rhabdomere microsaccades, aperture clipping, and the intracellular pupil). By providing a bridge between high-fidelity visual ecology and neuromorphic modelling, RhabdoForge enables researchers to explore how the interplay of sensory optics and neural processing can generate complex behaviour in both biological and artificial agents.

## 1 Introduction

Neuroethological models explore the neural mechanisms that underlie the real world behaviour of animals. Crucially, understanding behavioural capabilities cannot be based on modelling neural circuits in isolation, but needs to respect the following principles: 1-The evolution of the central neural hardware was shaped by the nature of the information present in the outside world. Each species’ own sensory and neuroanatomical “quirks” were directly influenced by the specific local challenges present in their respective ecological niches [1]; 2-Evolution has tuned the sensory systems to act as a pre-processors for the brain. Often, by its very geometry and/or physical properties, the sensory periphery filters and extracts the relevant information to be passed to the central brain [2, 3]; and 3-Information acquisition is a dynamic and closed-loop process: perception modulates the central brain, central brain drives action, which in turn alters perception. Thus, to truly understand behaviour, and replicate it through neuromorphic modelling, it is insufficient to model the brain alone [4, 5]. This is a multi-level challenge, and we must incorporate these other elements in our modelling efforts [6].

Visually-controlled behaviour in insects provides a relevant example. Insects use vision to solve complex real-world problems ranging from autonomous navigation in the desert to high-speed aerial pursuit with an efficiency that frequently rivals or exceeds modern engineering approaches [7–13]. They possess nervous systems that are orders of magnitude smaller than those of vertebrates, and are also much more energy-efficient [14–17]. This combination of minimal neural hardware and sophisticated behavioural repertoire makes insects an excellent model to study, both for understanding how neuroanatomy generates behaviour, but also as a source of inspiration for developing more efficient engineering solutions [18–20]. However, to date, models of these systems have largely used a ‘camera’ type eye for the input.

While the vertebrate eye with its single lens and retina is often viewed as the biological standard for visual sensing, it is an “evolutionary outlier”: the vast majority of visual systems on Earth are rhabdomere-based [21, 22]. Insects, in particular, have compound eyes consisting of thousands of individual optical units (ommatidia) with their lenses tiled as a curved surface, each containing several photoreceptor cells, with specialised structures called rhabdomeres, containing light-sensitive microvili [21, 23–25]. This architecture samples the world in a panoramic and highly parallel manner, which allows for several computational “shortcuts” to emerge directly from (or be greatly aided by) the physical organisation of the eye, including motion detection, low-pass filtering, and unique optic flow signatures [26–30]. More importantly, the compound eye is not a static collection of sensors: recent findings reveal that rhabdomeres within individual ommatidia move actively, a high-frequency movement that enables hyperacuity [31, 32].

For a neural model to provide an understanding of an insect’s vision-based behaviour, it should receive visual input that faithfully replicates these specific optics and internal dynamics. On the other hand, high-fidelity biophysical models such as [32, 33], are not practical to use in real-time, closed-loop simulations, and may be difficult to adapt to represent the eyes of different insect species. A promising approach to address these issues is provided by CompoundRay [34, 35], which models the multi-viewpoint nature of compound vision by leveraging hardware-accelerated ray-tracing [36, 37]. This approach essentially determines the radiance arriving at a given sensor location and orientation by ‘casting’ a ray from the sensor into the 3D (simulated) environment to intersect with surfaces, and thus inferring the property of the light reflected towards the sensor.

CompoundRay is currently the state-of-the-art for raw rendering performance of insect vision, capable of generating datasets of compound-eye views in arbitrary 3D environments at thousands of frames per second. However, this is obtained partly at the cost of simplifying the photoreceptor array within each ommatidia to a single unit, and perhaps more significantly, at the cost of reliance on proprietary hardware frameworks and low-level computational architectures that limits usability. This is particularly significant for neuromorphic modelling studies often involve an extensive exploratory phase for the researcher, where code goes through several iterations, conceptual hypotheses, and the ability to easily view how the (virtual) neurons react to different sensory inputs offers valuable insights to the researcher. Implementing novel biophysical features at the sensory periphery in CompoundRay can pose a significant engineering challenge for neuroscientists.

We have therefore developed RhabdoForge, a cross-platform, modular insect vision renderer designed specifically for ease of use, hardware agnosticism, and closed-loop interaction with neur-omorphic modelling research. Because most neuromorphic models are implemented in Python, RhabdoForge is built with a Python-centric philosophy, allowing researchers to integrate it with very little overhead into existing neuromorphic modelling frameworks (such as Brian2, Nengo, NEST or others) [38, 39]. By leveraging custom Python bindings to the TinyBVH library [40, 41], RhabdoForge achieves high-performance ray-casting (and efficient path-tracing) on any GPU architecture without the need for proprietary vendor-locked frameworks, and importantly introduces a fully parametrised ommatidial model that allows simulating any arbitrary compound eye with real-time sub-ommatidial rhabdomere actuation.

## 2 Methods

### 2.1 The rhabdomere as the fundamental sampling unit

The primary aim of RhabdoForge is to provide a fully open and accessible rendering and development environment for closed-loop neuromorphic research, where every biologically-relevant parameter is easily manipulatable. To support both simplified and biophysically rigorous models within a single framework, and contrary to previous engines that often treat the ommatidium as the unit viewpoint, RhabdoForge adopts a rhabdomere-centric model where individual rhabdomeres are the fundamental units of computation. Note that in this work, and following common usage, we use R1–R8 types to denote both the photoreceptor cells and their rhabdomeres (in other words, each cell is represented by its rhabdomere). In eyes with a fused rhabdom (where several cells share one waveguide), each cell is still represented by its own unit with its own spectral sensitivity: the units share a single optical axis (and therefore the same sampled rays), but the signal is returned per cell, so that spectral information is preserved and can be pooled or separated by downstream processing. Hereafter, *rhabdomere* refers to the model’s sampling unit, i.e. its unit of computation (position, optical axis, ASF) and *photoreceptor* to the cell (opsin, membrane, downstream wiring); see Table 1. In this architecture, a compound eye is represented as a massively parallel array of *M* rhabdomeres, where *M* = *N × R* (*N* ommatidia, each containing *R* rhabdomeres). Every rhabdomere is treated as an independent, active sampling unit with its own spatial coordinates, optical axis, angular sensitivity function (ASF), and internal state. Here, the ASF of a rhabdomere is the angular distribution of light it accepts (analogous to a point-spread function). This approach allows both “simplified” models (typically *R* = 1, a single unit representing either a fused rhabdom whose cells are not distinguished, or a single central rhabdomere like R7/8), as well as “full” models (e.g. *R* = 7 or 8 for *Drosophila*, [42]) to share the same underlying data structures and GPU kernels.

**Table 1:**
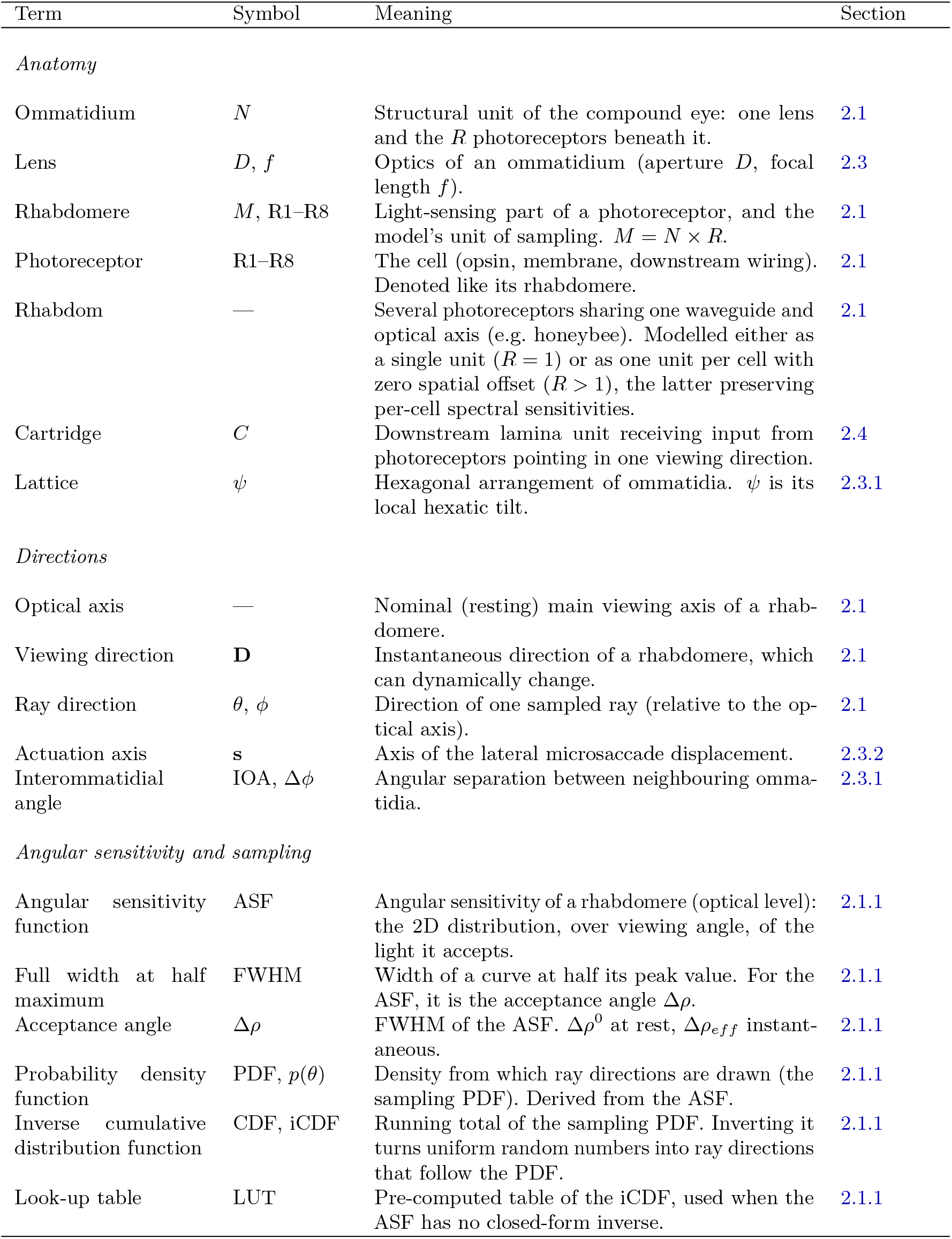
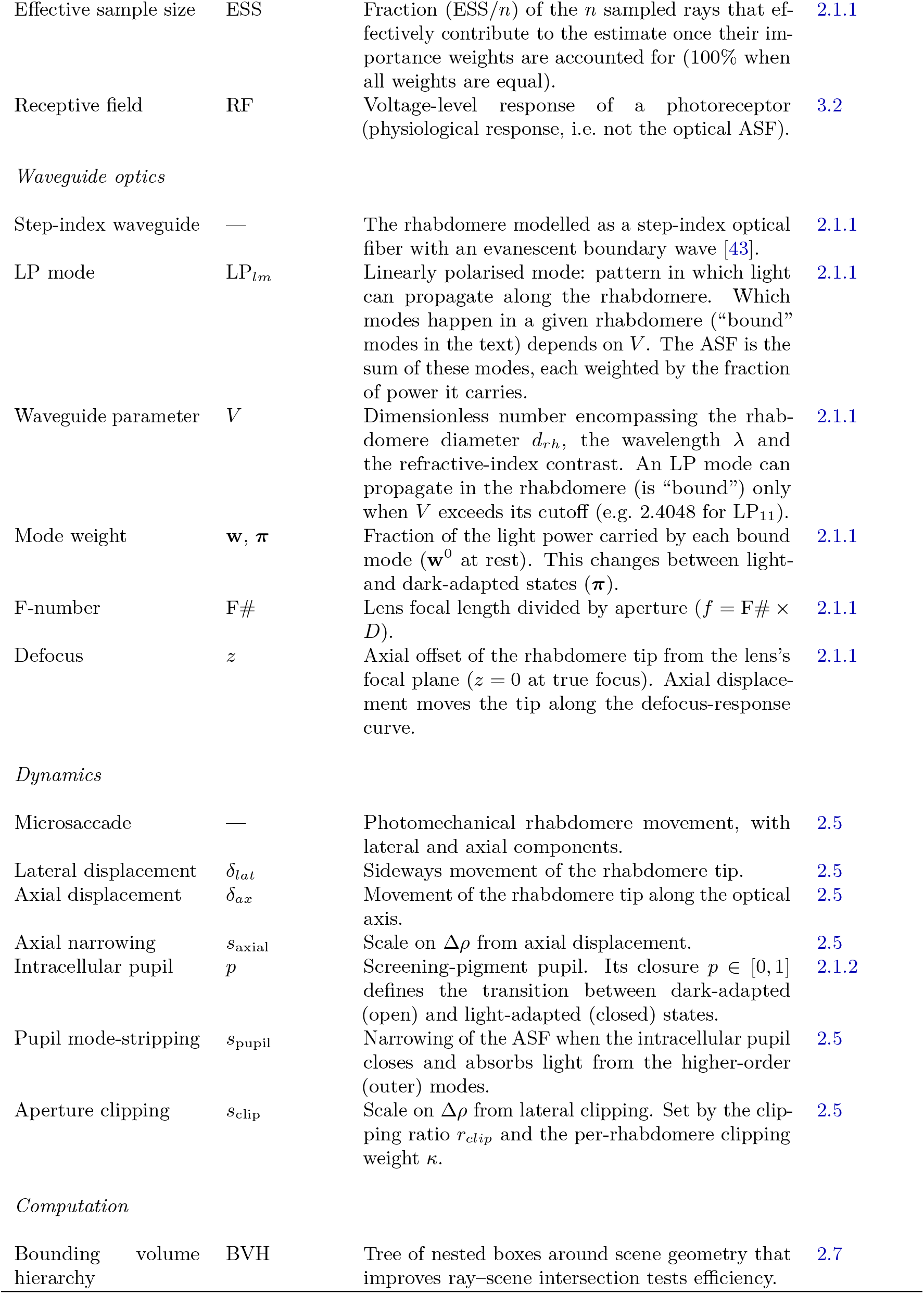
Terminology and notation used throughout the paper.

| Term | Symbol | Meaning | Section |
|---|---|---|---|
| *Anatomy* |  |  |  |
| Ommatidium | N | Structural unit of the compound eye: one lens and the R photoreceptors beneath it. | 2.1 |
| Lens | D, f | Optics of an ommatidium (aperture D, focal length f). | 2.3 |
| Rhabdomere | M, R1–R8 | Light-sensing part of a photoreceptor, and the model's unit of sampling. M = N × R. | 2.1 |
| Photoreceptor | R1–R8 | The cell (opsin, membrane, downstream wiring). Denoted like its rhabdomere. | 2.1 |
| Rhabdom | — | Several photoreceptors sharing one waveguide and optical axis (e.g. honeybee). Modelled either as a single unit (R = 1) or as one unit per cell with zero spatial offset (R > 1), the latter preserving per-cell spectral sensitivities. | 2.1 |
| Cartridge | C | Downstream lamina unit receiving input from photoreceptors pointing in one viewing direction. | 2.4 |
| Lattice | ψ | Hexagonal arrangement of ommatidia. ψ is its local hexatic tilt. | 2.3.1 |
| *Directions* |  |  |  |
| Optical axis | — | Nominal (resting) main viewing axis of a rhab-domere. | 2.1 |
| Viewing direction | **D** | Instantaneous direction of a rhabdomere, which can dynamically change. | 2.1 |
| Ray direction | θ, ϕ | Direction of one sampled ray (relative to the op-tical axis). | 2.1 |
| Actuation axis | **s** | Axis of the lateral microsaccade displacement. | 2.3.2 |
| Interommatidial angle | IOA, Δϕ | Angular separation between neighbouring omma-tidia. | 2.3.1 |
| *Angular sensitivity and sampling* |  |  |  |
| Angular sensitivity function | ASF | Angular sensitivity of a rhabdomere (optical level): the 2D distribution, over viewing angle, of the light it accepts. | 2.1.1 |
| Full width at half maximum | FWHM | Width of a curve at half its peak value. For the ASF, it is the acceptance angle Δρ. | 2.1.1 |
| Acceptance angle | Δρ | FWHM of the ASF. Δρ<sup>0</sup> at rest, Δρ<sub>eff</sub> instant-aneous. | 2.1.1 |
| Probability density function | PDF, p(θ) | Density from which ray directions are drawn (the sampling PDF). Derived from the ASF. | 2.1.1 |
| Inverse cumulative distribution function | CDF, iCDF | Running total of the sampling PDF. Inverting it turns uniform random numbers into ray directions that follow the PDF. | 2.1.1 |
| Look-up table | LUT | Pre-computed table of the iCDF, used when the ASF has no closed-form inverse. | 2.1.1 |
| Effective sample size | ESS | Fraction (ESS/n) of the n sampled rays that ef-fectively contribute to the estimate once their im-portance weights are accounted for (100% when all weights are equal). | 2.1.1 |
| Receptive field | RF | Voltage-level response of a photoreceptor (physiological response, i.e. not the optical ASF). | 3.2 |
| *Waveguide optics* |  |  |  |
| Step-index waveguide | — | The rhabdomere modelled as a step-index optical fiber with an evanescent boundary wave [43]. | 2.1.1 |
| LP mode | LP<sub>lm</sub> | Linearly polarised mode: pattern in which light can propagate along the rhabdomere. Which modes happen in a given rhabdomere ("bound" modes in the text) depends on V. The ASF is the sum of these modes, each weighted by the fraction of power it carries. | 2.1.1 |
| Waveguide parameter | V | Dimensionless number encompassing the rhab-domere diameter d<sub>rh</sub>, the wavelength λ and the refractive-index contrast. An LP mode can propagate in the rhabdomere (is "bound") only when V exceeds its cutoff (e.g. 2.4048 for LP<sub>11</sub>). | 2.1.1 |
| Mode weight | **w**, **π** | Fraction of the light power carried by each bound mode (**w**<sup>0</sup> at rest). This changes between light- and dark-adapted states (**π**). | 2.1.1 |
| F-number | F# | Lens focal length divided by aperture (f = F# × D). | 2.1.1 |
| Defocus | z | Axial offset of the rhabdomere tip from the lens's focal plane (z = 0 at true focus). Axial displace-ment moves the tip along the defocus-response curve. | 2.1.1 |
| *Dynamics* |  |  |  |
| Microsaccade | — | Photomechanical rhabdomere movement, with lateral and axial components. | 2.5 |
| Lateral displacement | δ<sub>lat</sub> | Sideways movement of the rhabdomere tip. | 2.5 |
| Axial displacement | δ<sub>ax</sub> | Movement of the rhabdomere tip along the optical axis. | 2.5 |
| Axial narrowing | s<sub>axial</sub> | Scale on Δρ from axial displacement. | 2.5 |
| Intracellular pupil | p | Screening-pigment pupil. Its closure p ∈ [0, 1] defines the transition between dark-adapted (open) and light-adapted (closed) states. | 2.1.2 |
| Pupil mode-stripping | s<sub>pupil</sub> | Narrowing of the ASF when the intracellular pupil closes and absorbs light from the higher-order (outer) modes. | 2.5 |
| Aperture clipping | s<sub>clip</sub> | Scale on Δρ from lateral clipping. Set by the clip-ping ratio r<sub>clip</sub> and the per-rhabdomere clipping weight κ. | 2.5 |
| *Computation* |  |  |  |
| Bounding volume hierarchy | BVH | Tree of nested boxes around scene geometry that improves ray–scene intersection tests efficiency. | 2.7 |

**Table 2:** Waveguide parameters and resting acceptance angles of our example *Drosophila* model. Reference lens (aperture 16.0 µm, F-number 1.288), dark-adapted, tip at the resting defocus *z* = *−*4.33 µm. *V* : waveguide parameter at the listed wavelength; Δ*ρ*: acceptance angle (FWHM).

| Cell | Diameter (µm) | Wavelength (nm) | *V* | Bound modes | Δ*ρ* (°) |
|---|---|---|---|---|---|
| R1 | 1.978 | 480 | 3.228 | 2 | 5.59 |
| R2 | 1.668 | 480 | 2.722 | 2 | 4.98 |
| R3 | 1.639 | 480 | 2.674 | 2 | 4.83 |
| R4 | 1.661 | 480 | 2.711 | 2 | 4.95 |
| R5 | 1.736 | 480 | 2.834 | 2 | 5.23 |
| R6 | 1.928 | 480 | 3.146 | 2 | 5.55 |
| R7/8 | 1.292 | 350 | 2.891 | 2 | 4.40 |

Each rhabdomere has a fixed location relative to its parent ommatidium (a focal-plane offset and optical axis). Throughout, the *optical axis* is the rhabdomere’s nominal (resting) axis, its *viewing direction* is its instantaneous direction (which can be changed by a microsaccade or a retinal shift), and each sampled ray has its own *ray direction*. How this layout is generated, and how it can depend on the ommatidium’s position in the array, is described later (see *Constructing an eye*). First we consider a single rhabdomere in isolation. In order to determine the light signal that each rhabdomere reports, we need to determine, for the model, the set of rays it casts into the 3D enviroment in order to render a given frame. This first requires a choice of optical model to determine the ASF, which sets the target probability density function (PDF) from which the ray directions are sampled. For rendering efficiency, it is desirable to use importance sampling directly from this PDF to cast the rays (rather than uniformly sampling all directions and then weighting each ray’s contribution by the ASF). This is however non-trivial for arbitrary ASFs, and we propose look-up table-based solutions for this (see *Custom ASFs and Exact iCDF LUTs* for static ASFs and *Mixture-of-Modes Importance Sampling* for the dynamic waveguide case). Finally, to draw random sample rays, there are several different randomness generators that can be used.

Each rhabdomere is thus processed in two decoupled GPU passes, linked by an intermediate buffer. In the *dispatch* pass, the rhabdomere casts *n* rays whose directions are drawn from its ASF (described below), and each ray writes its colour (three spectral channels) together with a scalar importance weight to the buffer. In the *reduction* pass, the contributions of these rays are accumulated into one value per spectral channel, i.e. three radiance values per rhabdomere. These accumulated values can then optionally go through optical attenuation, temporal filtering (membrane integration) and light adaptation (see *Reduction and adaptation*). Because the two passes only communicate through the buffer, they can use independent sampling and integration strategies.

#### 2.1.1 Angular sensitivity function (ASF)

To model the angular sensitivity of each rhabdomere, the ray casting engine stochastically generates sample ray directions whose spatial density matches that rhabdomere’s 2D ASF. RhabdoForge supports several optical models to define this ASF, which allow to scale from analytical approximations to detailed waveguide biophysics. The sampling strategies described below only determine how a given ASF is evaluated by Monte Carlo integration. They all converge on the same ASF, and differ in noise and computational cost, not in the modelled optics.

##### Analytical Models and Snyder Optics

The baseline ASF is an anisotropic Gaussian (a tilted ellipse parameterised by acceptance angles Δ*ρ*_*maj*_ and Δ*ρ*_*min*_, each defined as a Full Width at Half Maximum, FWHM). The acceptance angles (Δ*ρ*) can be static or dynamically (see *Sub-ommatidial biophysics and dynamic optics*) driven by Snyder’s optical formulation [44], which approximates the rhabdomere as a sharp geometric cone and combines geometric blur with the wavelength-dependent lens diffraction limit.

##### Waveguide Optics

For a more realistic and non-Gaussian ASF, the engine also implements waveguide optics following [43, 45]. Rather than a geometric cone and a point focus, the rhabdomere is modelled as a step-index optical fiber with an evanescent boundary wave, while the lens (operating at low Fresnel numbers) generates an elongated 3D diffraction volume. The engine solves for the bound linearly polarised (LP) waveguide modes using Bessel-function zeros. The rhabdomere’s overall ASF is the sum of these bound modes, weighted by their in-boundary power fractions. The acceptance angle Δ*ρ* is the FWHM of this summed ASF. For multi-mode rhabdomeres it is measured outward from the ASF’s peak, to account for modes that peak off-axis.

Crucially, this excitation integral includes a defocus term (*z*). In species such as *Drosophila*, the rhabdomere tip rests several micrometres short of the true focal plane (17.0 µm behind the lens, while the back focal plane lies at 21.33 µm, placing the tip at a resting defocus of *z* = *−*4.33 µm [32]). Accounting for this resting defocus is important: at true focus (*z* = 0), waveguide optics offer no clear difference over Snyder’s formulation, but a defocus of *z* = *−*4.33 µm places the rhabdomere on the steep flank of the defocus-response curve, meaning any axial photomechanical movement would thus dynamically alter the acceptance angle (see *Sub-ommatidial biophysics and dynamic optics*) in a way that plain geometric optics cannot reproduce.

##### Exact Importance Sampling

The simplest way to integrate over an ASF is to cast rays uniformly over directions and weight each one by the ASF. This however means that many rays will carry very little weight (in the tails), and are thus essentially wasted computation. An alternative is to draw ray directions directly from the ASF using pure importance sampling [46, 47], so that the density of rays already embeds the ASF and all rays can be weighted equally. For the Gaussian ASF, given a uniform random variable *U*_1_ ∈ [0, 1), the corresponding sample’s radial deviation *θ* from the optical axis for the major and minor axes is generated using the inverse Cumulative Distribution Function (CDF):

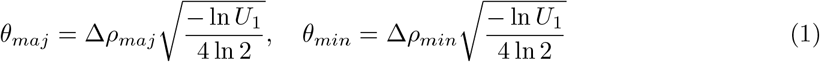

The constant 4 ln 2 ≈ 2.77 ensures that Δ*ρ* corresponds to the FWHM. Azimuthal distribution around the optical axis is handled by a second uniform variable *U*_2_ ∈ [0, 1) to generate an angle *φ* = 2*πU*_2_. These sampled coordinates are then rotated by the local hexatic tilt *ψ*_*i*_ of the hexagonal ommatidial lattice and mapped to a local tangent plane before transformation into world coordinates. This is, as far as the literature describes, the scheme used by existing compound-eye renderers [34, 35]. It is efficient, but requires the ASF to have an analytically invertible cumulative distribution, which in practice restricts it to a Gaussian.

##### Custom ASFs and Exact iCDF LUTs

For arbitrary (non-Gaussian) static ASFs (e.g. diffraction-limited Airy disks), analytical inversion of the radial ASF’s cumulative distribution is impossible. RhabdoForge thus computes a discretised, area-weighted inverse Cumulative Distribution Function Look-Up Table (iCDF LUT) at eye construction. The radial ASF is discretised into a piecewise-linear table of length *L* = 256 over a radius range of 4*×* FWHM. On the GPU, the sample rays are then drawn via exact importance sampling by looking up uniform random variables *U*_1_ directly in this iCDF LUT. Discretisation across *L* = 256 bins keeps radial quantisation errors (*∼*1.3 *×* 10^*−*4^ FWHM) well beneath stochastic Monte Carlo noise, while the truncation boundary at 4 *×* FWHM captures *>* 99.99 % of the optical power for Gaussian and waveguide ASFs. Because the LUT only needs the radial ASF to be tabulated, the same procedure applies to any other static, user-supplied ASF (such as the Airy disk mentioned above). It is worth noting that the cumulative distribution flattens towards 1.0, so the last bin of the LUT spans a large radius range: the outermost *∼* 0.4 % of the probability mass are sampled with a coarser, one-sided approximation. This however stays (at the ray counts typically used) below the Monte Carlo noise. Second, the 4*×* FWHM truncation is a per-ASF assumption: it costs nothing for a Gaussian or a waveguide ASF, but a heavy-tailed custom ASF (e.g. Cauchy-like) would lose probability mass (and bias the estimate low) in the tails.

##### Mixture-of-Modes Importance Sampling

With waveguide optics, the ASF changes shape non-linearly as intracellular pupil closure and axial microsaccades modulate the multi-mode power balance in the rhabdomere waveguide [43, 48], (see *Reduction and adaptation*). Recomputing and re-inverting an exact iCDF on the host CPU or GPU for every rhabdomere at every frame is computationally prohibitive. Therefore, when waveguide optics are active, the engine introduces a mixture-of-modes importance sampler: at eye build time, the engine pre-solves the continuous ASF of each individual bound LP mode across an 8 *×* 8 grid of pupil and axial-displacement states, and this grid is then baked into a 2D texture. The intracellular pupil absorbs evanescent power without reshaping a mode’s ASF, and so the pupil-closure axis only acts as a pure rescaling of the mode weights, while the rhabdomeres axial displacement alter modes geometry.

At runtime, the GPU interpolates the current weight and width of each bound mode. The sampler probabilistically selects a mode based on its weight, draws a sample from a proposal PDF (a broadened Gaussian, *σ* = 1.7*×* current mode width), and applies an exact unbiased reweighting against a bilinear texture lookup of the true summed-mode ASF.

##### Anatomical inputs

For the *Drosophila* eye model used in our examples, anatomical parameters were derived from empirical reconstructions: an object focal length of 20.6 µm (computed from the thick-lens parameters of [43], who reports 20 *±* 2 *µ*m) and a reference lens aperture of 16.0 µm (F-number 1.288). Individual rhabdomere diameters are specified per cell (R1–R6: 1.978 µm, 1.668 µm, 1.639 µm, 1.661 µm, 1.736 µm and 1.928 µm; R7/8: 1.292 µm; redigitised from [31]). R1–R6 rhabdomeres are evaluated at 480 nm, making their waveguide parameter *V* ≈ 2.67–3.23 above the LP_11_ cutoff (2.4048) so that all outer rhabdomeres display two bound modes. At 1.292 µm and its UV opsin peak (350 nm), R7/8 also displays two bound modes (*V* = 2.891). Note that R7/8 is modelled as a single central unit: in the fly, R7 (UV-sensitive) and R8 (blue/green-sensitive) are stacked above each other along the optical axis.

To make rhabdomere bundle blueprints interchangeable across arbitrary eye geometries, the rhabdomere bundle defines the F-number (1.288), while individual lenses provide physical apertures. Lens focal lengths are computed as *f* = F#*×* aperture, and the resting defocus scales proportionally with focal length. Waveguide parameters that depend on local lens geometry are pre-solved on a 6 *×* 6 interpolation grid over the eye’s F-number and focal length range, and interpolated per lens at eye construction.

To draw random ray directions from the PDF described above, RhabdoForge provides several interchangeable generators that balance stochasticity, hardware performance, and realism (see *Random generators* suplemental material section). All the generators can be combined with temporal dithering, where a per-frame counter perturbs the sample stream so that the exact ray directions fluctuate stochastically from one time step to the next. The temporal stochasticity that is introduced can then be accumulated by temporal integration (see below), or by downstream neuromorphic circuits, and helps mitigate structured spatial aliasing (Moiré patterns caused by high-frequency spatial structures, [49], or the residual noise of path-tracing mode).

Together, the ASF, the sampling strategy, and the chosen random generator fully determine the set of rays a given rhabdomere casts in a given frame. The reduction pass that accumulates the contributions of these rays is described in the next section.

#### 2.1.2 Reduction and adaptation

Once the rays for a rhabdomere have been cast in the dispatch pass, a GPU reduction pass integrates the *n* per-ray contributions of the current frame into a raw signal. This is computed as a weighted mean, *Ĉ* = ∑_*i*_ *w*_*i*_*C*_*i*_ */* ∑_*i*_ *w*_*i*_, where *C*_*i*_ and *w*_*i*_ are the colour and importance weight written by ray *i* (under pure-importance sampling, all weights are 1.0 and this collapses to a simple arithmetic mean).

The GPU pipeline applies optical attenuation, temporal filtering, and light adaptation before returning the data to the downstream neuromorphic model on the CPU. For each frame, the data returned is a four-component vector per rhabdomere: the first three values represent the temporally integrated and gain-controlled responses across three discrete spectral channels. These channels can be evaluated at user-defined wavelengths (by default approximating display primaries at 610 nm, 550 nm, and 465 nm) weighted against an empirical opsin template (the Govardovskii A1 template [50]) at the photoreceptor’s peak wavelength, evaluated at a single wavelength per channel. The fourth value reports the instantaneous biochemical gain factor. Providing this raw gain factor in the fourth channel allows to “un-bake” the adaptation and decouple the stages if desired.

##### Optical attenuation (Intracellular pupil)

In addition to biochemical adaptation, Rhab-doForge models the intracellular screening-pigment pupil [48, 51]. In vivo, dark pigment granules migrate toward the rhabdomere boundary upon light exposure, absorbing the evanescent wave of bound waveguide modes. This mechanism is driven by a dedicated, seconds-scale exponential moving average (EMA) (e.g. *τ*_*pupil*_*≈* 2.0 s, within the 1 s to 10 s dynamics reported in [31]), producing two distinct physical effects:

1. **Spatial narrowing**: Higher-order waveguide modes are selectively stripped, narrowing the resting acceptance angle Δ*ρ* to 55 % to 65 % of its dark-adapted width in R1–R6 and 55 % in R7/8 (see *Sub-ommatidial biophysics and dynamic optics*), close to the factor of *∼*0.6 obtained for the fully closed pupil in [51].
2. **Optical transmittance reduction**: Stripping modal power induces a direct 4–12*×* reduction in transmittance at full closure (transmittance drops to 0.08–0.22 in R1–R6 and 0.25 in R7/8), in line with the decrease in absorption of up to six-fold estimated for the housefly [51].

Because the waveguide parameter *V* is inversely proportional to wavelength (*V ∝* 1*/λ*), this transmittance loss is computed independently per spectral channel (whereas the narrowing ratio is solved once per cell, at its own peak wavelength), and applied during the reduction pass, attenuating the raw radiance before it reaches subsequent biochemical adaptation stages. Non-waveguide optical models bypass this mechanism and keep transmittance at unity.

Concretely, each ommatidium carries a pupil closure *p ∈* [0, 1] (0: dark-adapted, 1: fully closed) that relaxes with time constant *τ*_*pupil*_ towards a saturating function of the slow luminance tracker (defined below), *p*_*∞*_ = *Ī/*(*Ī* + *I*_*ref*_ ), so that the pupil is half closed at the reference luminance *I*_*ref*_ . The narrowing and transmittance factors are pre-solved for the fully closed state (per cell, and per spectral channel for transmittance) and interpolated linearly with *p* between their dark-adapted value of 1 and the closed value: *s*_pupil_ = 1 *−p* (1 *−s*_closed_) (and similarly for the transmittance).

##### Luminance tracking and biochemical gain control

The engine maintains two exponential moving averages (EMAs) of incident radiance at the ommatidium level: a *fast* tracker (*Î* with *τ ≈* 5 ms for *Drosophila*) and a *slow* tracker (*Ī* with *τ ≈* 100 ms) which phenomenologically reflect the timescales of PIP_2_ hydrolysis and Ca^2+^ dynamics, respectively [31, 52]. The slow tracker drives a hyperbolic (Naka-Rushton-type, with unit exponent) divisive adaptation gain [53]:

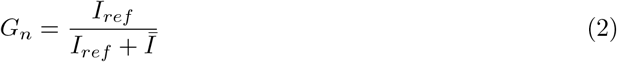

which scales the response gain based on the local operating luminance, compressing high-dynamic-range scenes into the operating range of the simulated photoreceptors [54, 55].

##### Membrane integration

The resulting signal can be subjected to a final temporal low-pass filter representing the RC integration time of the photoreceptor membrane (*τ*_*membrane*_). Given a time constant *τ*_*r*_ (around 10 ms to 15 ms in *Drosophila* R1–R6 cells, [56]) and the wall-clock inter-frame interval Δ*t*, the filtered output 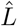 is updated each frame as:

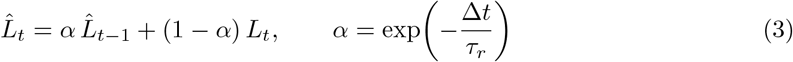

where *L*_*t*_ is the current frame’s adapted radiance. This GPU-side membrane low-pass filter serves as a minimal but biophysically grounded preprocessing stage. The full phototransduction cascade involves, in reality, many additional non-linear dynamics [33, 57]. Disabling this filter simply returns the instantaneous mean radiance, which can therefore be coupled to arbitrarily detailed downstream biophysical models of photoreceptor responses on the Python side if desired.

The choice of membrane integration time, relative to the frame interval, will affect the optimal choice of sampling method (see above, and *Random generators* supplemental material section). That is, because this process integrates each rhabdomere’s samples over time, the effective sampler quality is set not by single-frame discrepancy but by behaviour under temporal accumulation, attenuating the frame-to-frame Monte Carlo variance introduced by temporal dithering. Progressive sequences whose index advances each frame (see *Random generators* supplemental material section), such as Halton or Sobol, keep refining the estimate as frames accumulate, whereas fixed-*n* constructions (those that are only re-randomised per frame: Hammersley, Fibonacci, stratified) only average down as independent samples. Hence, the longer a rhabdomere integrates, the more a progressive sampler will outperform a lower-discrepancy (but non-extensible) one.

### 2.2 From rhabdomeres to downstream models

These filtered and gain-controlled per-rhabdomere responses are the engine’s main output: an array of *M* = *N × R* rhabdomeres, each with three spectral channels plus the adaptation gain. Because RhabdoForge does not aim to model what happens downstream of the photoreceptors, this array can be passed to any neuromorphic model, which is thus free to combine it as it requires (e.g. **c**(*t*) = **W r**(*t*) for a linear mapping **W** from rhabdomeres to lamina units).

RhabdoForge does however provide the anatomical wiring that such a model needs: for each cartridge, which are the rhabdomeres from neighbouring ommatidia that project to it. Indeed, in neural superposition eyes, the outer photoreceptors R1–R6 of neighbouring ommatidia that share a viewing direction converge onto the same cartridge, whereas the central photoreceptors (the merged R7/8 in our example *Drosophila* bundle, which projects to the medulla) are not pooled across ommatidia and project directly. The specific structure of this wiring and how the engine derives it are detailed later (see *Cartridge wiring: simple versus neural superposition* and the Results). Note that we distinguish the cartridge from the laminar cells within it (e.g. L1–L3 in *Drosophila*): the wiring only specifies which photoreceptors feed a given cartridge, not how each laminar cell weights them.

In apposition eyes with fused rhabdoms (such as the honeybee), the rhabdomeres share a single optical axis with essentially zero spatial offset. The scene is therefore sampled once per ommatidium, and each photoreceptor’s own spectral sensitivity is then applied to that radiance, meaning that output remains a four-component vector per receptor (preserving the spectral channels). Note that the renders in Fig. 6 use one pooled value per ommatidium and per colour, purely for display.

On the Python side, these raw arrays are mapped to a GPU-ready buffer and exposed (wrapped) through multiple “proxies”, object-oriented views which allow fast, NumPy-like slicing [58] and vectorised mathematical operations natively in Python. This abstraction layer allows researchers to query the output using intuitive biological pathway concepts (e.g. output from all rhabdomeres that correspond to the motion detection pathway, or from rhabdomeres that correspond to the colour vision), by physical structures (e.g. per ommatidium, per cartridge), or per property (e.g. per colour channel), reducing the boilerplate code required to interface with downstream neuromorphic frameworks.

### 2.3 Constructing an eye

An eye is represented as a curved 3D lattice of ommatidia. Each ommatidium carries a set of rhabdomeres with a specific layout relative to its optical axis, and that layout can itself depend on the ommatidium’s location within the array (e.g. the equatorial chirality flip described below). Building an eye therefore means (i) generating the hexagonal lattice of ommatidial lenses, (ii) assigning each ommatidium its internal rhabdomere layout, and (iii) deriving the per-ommatidium and per-rhabdomere optical properties.

A core architectural philosophy of RhabdoForge is the decoupling of the sensory “layers”. Rather than treating a compound eye as a monolithic asset, the engine treats the global shell (the eye 3D shape), the lattice topology (lens distribution on that surface), and the rhabdomere blueprint (internal rhabdomere organisation, along with their photomechanical fields) as orthogonal components. This modularity allows to construct high-fidelity replications of known morphologies or, alternatively, to create “chimeric” eyes (for example, mapping a *Drosophila*-style rhabdomere bundle onto a honeybee ommatidium lattice).

A recurring hurdle in compound vision simulation is the acquisition of anatomically accurate eye models. While high-resolution micro-CT scans of insect heads, and computational tools to reconstruct accurate 3D models, are becoming more common [59–66], they remain largely inaccessible for many non-model species. Consequently, many compound eye simulations rely on simpler approximations (i.e. uniform spherical models). To address this gap and provide immediate utility to the neuromorphic community, RhabdoForge provides a small but extensible morphological library of biologically-seeded procedural generators alongside a flexible pipeline for reconstructing eyes from sparse empirical data.

#### 2.3.1 Generating the lattice

We provide native reimplementations of procedural ommatidial lattice generation algorithms from the literature. This includes an adaptation of the Stürzl honeybee (*Apis mellifera*) eye model [67], which defines four distinct zone boundaries and empirical formulas for horizontal and vertical interommatidial angles (IOA), as well as the *Drosophila melanogaster* model used by [32], which simulates lattice growth via a Breadth-First Search (BFS) algorithm across a constrained 3D morphology.

However, much of the empirical data regarding insect eye resolution was recorded before the advent of 3D micro-CT scanning, often published as 2D stereographic projections or Mercator maps of ommatidial viewing directions (e.g. [24, 68]). To take advantage of this vast anatomical literature, RhabdoForge includes an optimisation pipeline that maps these 2D records onto fully functional 3D sensory arrays.

Provided 2D coordinates (for instance by digitising such a 2D plot in SVG), the module estimates two continuous fields by Radial Basis Function (RBF) interpolation: a local spacing field (the reciprocal of the ommatidial density *ρ*(**x**)), and a local hexatic-axis field *ψ*(**x**) recovered from the Ψ_6_ phase of the source coordinates. A theoretical hexagonal lattice, whose unit-cell angles are themselves measured from the traced source rows, is then clipped to the eye contour and refined through four stages:

(i)A global *density warp* displaces the clipped lattice radially to match the target spacing field. *(ii)* An *orientation-aware spring relaxation* treats the Delaunay triangulation as a network of springs whose rest lengths match the local spacing field, with edge directions snapped toward the nearest lattice bond to curve rows along the measured grain. Virtual points outside the boundary exert outward pressure, preventing edge collapse. *(iii)* A *density-correcting transport* pass solves a linearised optimal-transport Poisson equation via FFT to eliminate residual density errors without perturbing the local orientation. *(iv)* A *boundary-finalisation* stage culls out-of-domain points, merges duplicates, seeds new points along the smooth contour wherever a large gap remains, and settles the boundary with a short relaxation pass.

##### 3D surface reconstruction from normals

These unprojected 2D viewing directions, treated as the target optical axes, correspond to surface normals on a curved surface (ellipsoid slice). The algorithm solves for 3D lens coordinates via least-squares optimisation over the first-ring Delaunay edges:

1. Each edge chord length is constrained to match the prescribed physical lens diameter.
2. Each chord is constrained to be strictly perpendicular to the average of its two terminal optical axes.
3. A weak anchor to an empirical bounding volume (e.g. an ellipsoid, see below) resolves translation and global scale.

Under this formulation, the target lens size is thus explicitly maintained, while the local radius of curvature emerges naturally as the ratio of lens spacing to IOA (*R* ≈ spacing*/*IOA, median 178 µm). The resulting mesh exhibits excellent surface tangency (median chord-to-normal error of only 1.2^*°*^).

We provide an anatomically grounded *Drosophila* model using an ellipsoid curvature model derived from *Drosophila* head proportions (roughly 24 % taller than deep [69]), incorporating an anterior-ventral lens-size expansion gradient [70] with a median target lens diameter of *d* ≈ 17.0 *µ*m. Apertures are estimated using local topological spacing (median distance to first-ring neighbours). The resulting digital scaffold yields a median lens aperture of 16.78 µm (5th–95th percentile 14.9 µm to 18.5 µm), closely matching empirical measurements in *Drosophila melanogaster* (16.85(21) µm [71]). This data-driven pipeline allows researchers to rapidly generate anatomically faithful compound eyes for novel species simply by supplying a digitised 2D plot from existing literature.

#### 2.3.2 Default rhabdomere layout: feature derivation and modular overrides

Once the lens lattice is established, RhabdoForge allows for independent injection of ommatidium- or rhabdomere-level features. If specific parameters (such as IOAs or rhabdomere bundle orientation) are not manually provided from empirical data, the system implements a hierarchical “feature derivation” pipeline that can extract them from the local geometry.

For instance, the local lattice orientation for each ommatidium *j* is determined using the hexatic order parameter (Ψ_6_) [72]. We project the relative positions of the nearest neighbours *N*_*j*_ into the local tangent plane and derive the structural orientation *ψ*_*j*_ from the phase of the average hexatic phasor:

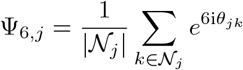

The magnitude |Ψ_6,*j*_| serves as a local metric of lattice regularity, while the orientation is recovered as *ψ*_*j*_ = ^1^ arg(Ψ_6,*j*_). The IOAs Δ*φ* are calculated by averaging the angular separation of neighbours along the primary lattice axes. For the simple sampling-based acceptance model, these values are used to scale the rhabdomere acceptance angles Δ*ρ* via the eye parameter *p* (Δ*ρ* = *p* Δ*φ*) [73]. For waveguide optics, by contrast, Δ*ρ* is set by the rhabdomere diameter, the wavelength and the lens focal length, and the interommatidial angle does not enter.

Crucially, these derivation rules are modular and can be selectively overridden. One can “mix and match” these components, for example keeping a biologically-derived ommatidial lattice but enforcing acceptance angles from a much lower-resolution eye, or orienting rhabdomere bundles to match a hypothetical optic flow field. This enables a combinatorial exploration of the design space of compound vision otherwise inaccessible with static eye models.

#### 2.3.3 Specific layout example: orienting the Drosophila rhabdomere bundle

In many insect eyes, individual ommatidia contain multiple rhabdomeres (for instance *R* = 8 in *Drosophila*) each with their distinct optical axes [42]. Signals from adjacent ommatidia that share the same visual axis converge onto a single neural relay in the Lamina, a principle known as neural superposition [74, 75]. In other words, two rhabdomeres in the same ommatidium “see” slightly different points in space, and conversely, two rhabdomeres from two neighbouring ommatidia can have the same viewing direction. The specific rhabdomeres that converge onto a single downstream neural relay, a so-called “cartridge”, therefore follow a very specific wiring [76–80].

It has been shown in *Drosophila* that the orientation of these rhabdomere bundles is remarkably precise and ecologically tuned: the main structural axis of the rhabdomere bundle is maintained, on average, at an 81^*°*^ offset relative to the projected anterior-posterior optic flow (accounting for the typical *∼* 10^*°*^ head pitch observed during flight) [32]. Importantly, this organisation displays an “equatorial shift”, where the rhabdomere bundle’s chirality is mirrored across the eye’s equatorial plane [81].

*In vivo*, this precise alignment and its downstream neural wiring are emergent outcomes of complex developmental cascades governed by planar cell polarity (PCP) signaling, tissue mechanics, and axonal self-sorting during pupal morphogenesis [76, 77]. Because simulating these biological growth dynamics is computationally prohibitive, RhabdoForge adopts an inverse, top-down approach: we take the empirical 81^*°*^ flight-aligned invariant as a macroscopic guiding target to generate an initial, smooth vector field across the lattice.

This orientation field serves a dual purpose: it establishes the anatomical rhabdomere yaw (*χ*_*i*_) and provides the structural prior required to solve the neural superposition wiring. On curved or irregular lattices, identifying which neighbouring ommatidia share a viewing direction cannot be solved by naive angular searches (e.g. greedily linking ommatidia whose optical axes are most nearly parallel), as local packing defects would lead to topological tearing and biologically impossible connections. Instead, RhabdoForge uses the flow-aligned bundle orientation as a structural template to select candidate donor ommatidia, unifies them into a globally valid wiring topology via Mixed-Integer Linear Programming (MILP), and retroactively fine-tunes each bundle’s geometric yaw to match the selected optical axes (see *Cartridge wiring: simple versus neural superposition*).

To construct this initial orientation field across procedurally generated or reconstructed eye meshes, our algorithm first calculates an alignment phasor field from a global optic flow pole (e.g. forward flight) as an alignment phasor field. This phasor field is then smoothed using a nematic (vectors treated as line segments, where *v* and *−v* are equivalent, to avoid issues with 180^*°*^ phase flips in the orientation field) k-nearest-neighbour (kNN) pass, on a strict per-chirality-zone fashion.

A standard projection of optic flow suffers from a singularity at the anterior pole (the point of expansion), where the flow magnitude vanishes and directions become radial. To replicate the biological organisation, which does not show such radial expansion field, our algorithm performs a “combing” effect near the expansion point, where the raw flow projection is blended with a chirality-zone-specific diagonal pull. This ensures that rhabdomere bundles in the frontal eye regions do not radiate from a single point, but instead maintain the oblique, coherent orientations observed in-vivo [32]. This results in a piecewise-smooth vector field in each hemisphere of each eye (for a target angle of 81^*°*^, mean = 81.1^*°*^, std = 2.3^*°*^; 78 % within *±* 2.0^*°*^), closely matching the biological distributions measured in *Drosophila* [32] and preserving the characteristic equatorial discontinuity (sharp “chevron-like” or “V-shape” kink at the equator where the hemispheric chirality and orientation flip), where deviations from pure flow are most pronounced in a “X” pattern in the frontal region (Fig. 4).

**Figure 1:**
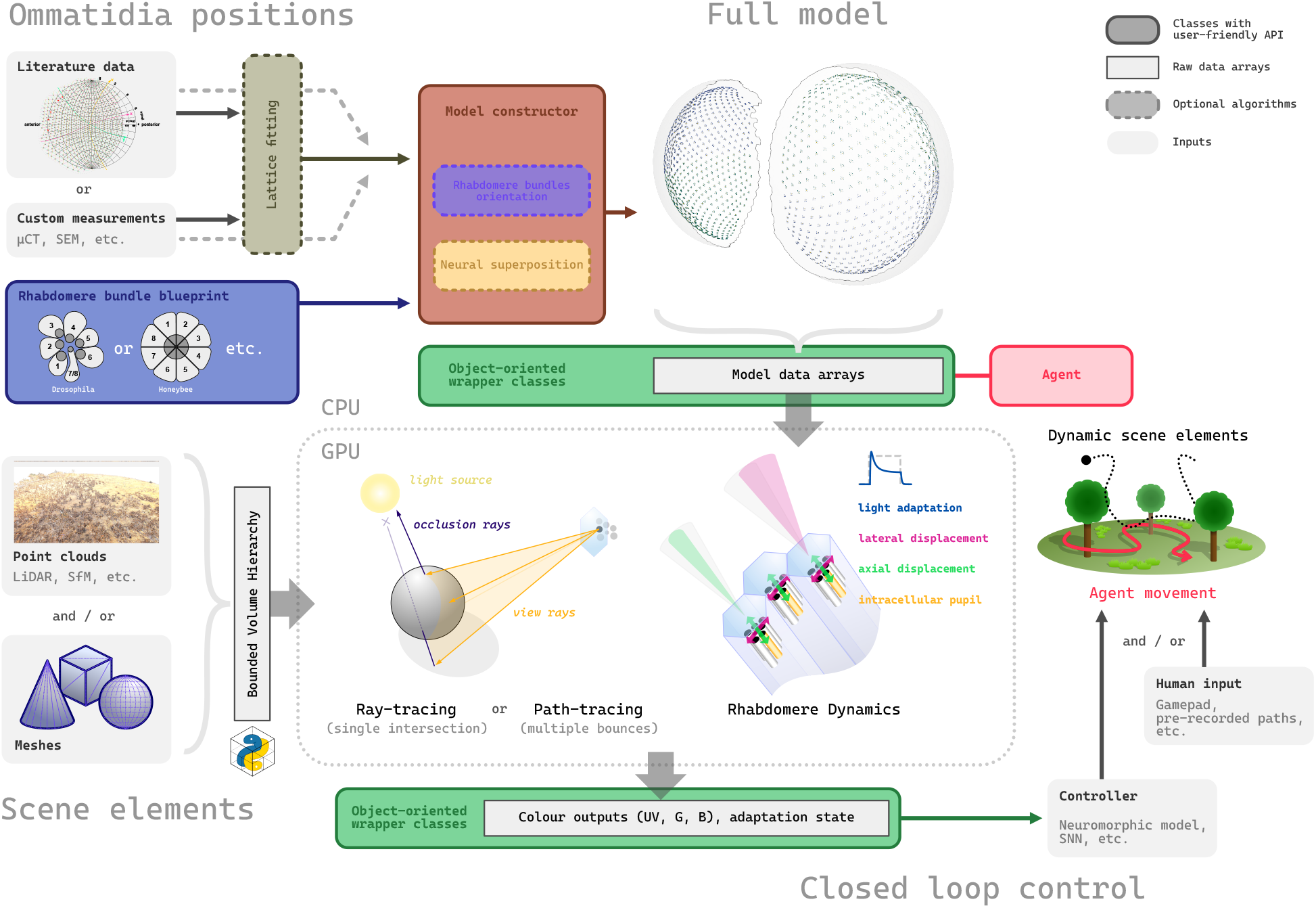
Overview of the RhabdoForge rendering architecture and closed-loop pipeline. Eye model construction and morphological pipeline. Compound eye geometries can be imported from empirical sources (literature-derived 2D projections, *µ*CT, SEM) or procedurally generated. An automated *Lattice fitting* stage reconstructs the 3D ommatidial array. Discrete *Rhabdomere bundle blueprints* (e.g. fused rhabdom such as honeybee or multi-rhabdomere such as *Drosophila*) are assigned to each ommatidium. The model constructor then computes optic-flow-aligned bundle orientations and, optionally, resolves downstream *Neural superposition* wiring. The resulting data structures are uploaded to GPU memory, and can be accessed and interacted with on the CPU side via Python wrapper proxies. Environments composed of polygon meshes and/or dense point clouds (e.g. LiDAR scans, photogrammetry) are partitioned into static (BLAS) and dynamic (TLAS) Bounding Volume Hierarchies via PyTinyBVH [40] for high-throughput ray-scene intersection tests. RhabdoForge supports both single-hit ray-tracing and multi-bounce stochastic path-tracing using exact importance sampling, iCDF LUTs or a mixture-of-modes importance sampling strategy. At each frame, dynamic GPU compute passes simulate sub-ommatidial biophysics in real-time, including local light adaptation (fast/slow luminance tracking, and intracellular pupil adaptation), retinal sheet movement, lateral and axial photomechanical microsaccades, and dynamic waveguide optics. Integrated per-photoreceptor spectral outputs (customisable to user-defined display-primary and photoreceptor peak wavelengths) and instantaneous adaptation states are read back to the CPU with minimal latency. Downstream controllers (e.g. Spiking Neural Networks, neuromorphic models) or external human/trajectory inputs modulate agent kinematics and dynamic scene elements, closing the real-time sensorimotor loop.

**Figure 2:**
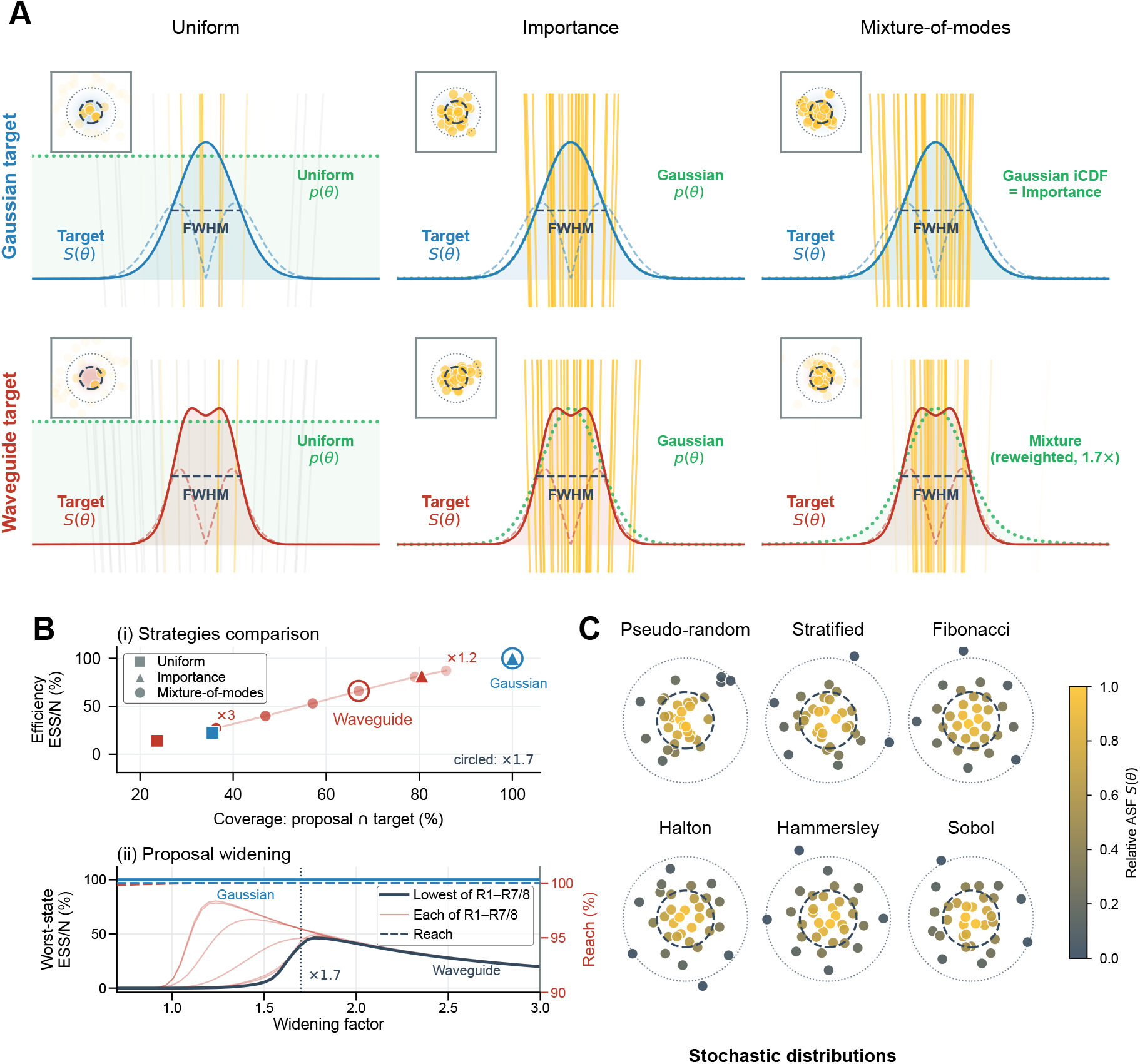
Importance sampling strategies and stochastic sample distributions. **(A)** Comparison of Monte Carlo integration strategies (Uniform, Importance, Mixture-of-modes) for a pure Gaussian (solid blue) and a biophysical Waveguide (solid red, showing the resting R1 ASF) target ASF. Dotted green: proposal PDF *p*(*θ*). Thin dashed curves: area-weighted ASF *S*(*θ*) *· θ*. Rays (*n* = 32) are weighted by opacity (grey: negligible weight). *Uniform*: evenly drawn samples (inefficient). *Importance*: exact for the Gaussian target ASF (all unit weights), but undersamples the tails of more complex ASFs. *Mixture-of-modes*: picks a specific waveguide mode by its live weight, draws from a broadened Gaussian proposal (*σ* = 1.7*×* ), and reweights against the true ASF. (Note: For the Gaussian target ASF, the Mixture-of-modes strategy corresponds to exact importance sampling). **(B)** Sampling efficiency, measured by the effective sample size (ESS*/n*): the fraction of the *n* rays that effectively contribute to the estimate once their importance weights are accounted for (100 % when all weights are equal, lower when few rays carry most of the weight). *(i)* Efficiency vs. Coverage (overlapping coefficient, 1 *−* TVD). Marker shape: strategy; colour: target ASF. For the Mixture-of-modes strategy, the circled marker denotes the chosen 1.7*×* widening factor. *(ii)* Proposal widening factor versus worst-case efficiency (solid, left axis) and signal reach (dashed, right axis). The Gaussian reference is invariant at 100 %. For the Waveguide target ASF, the widening factor is swept across all 7 rhabdomere types, each evaluated over its full biophysical grid of pupil-closure and axial-displacement states. Thin red lines show the worst state for each type; the bold black line shows the global worst-case envelope. The selected 1.7*×* widening factor allows Monte Carlo integration across all dynamic states. **(C)** Point set generators (*n* = 32) mapped to disk coordinates (*u*_1_, *u*_2_), coloured by the relative ASF value *S*(*θ*) (yellow: peak signal; dark: peripheral). Low-discrepancy sequences (Halton, Sobol, Hammersley) and geometric sets (Stratified, Fibonacci) prevent sample clumping seen in pseudo-random noise. Dashed circles denote FWHM and main lobe limits.

**Figure 3:**
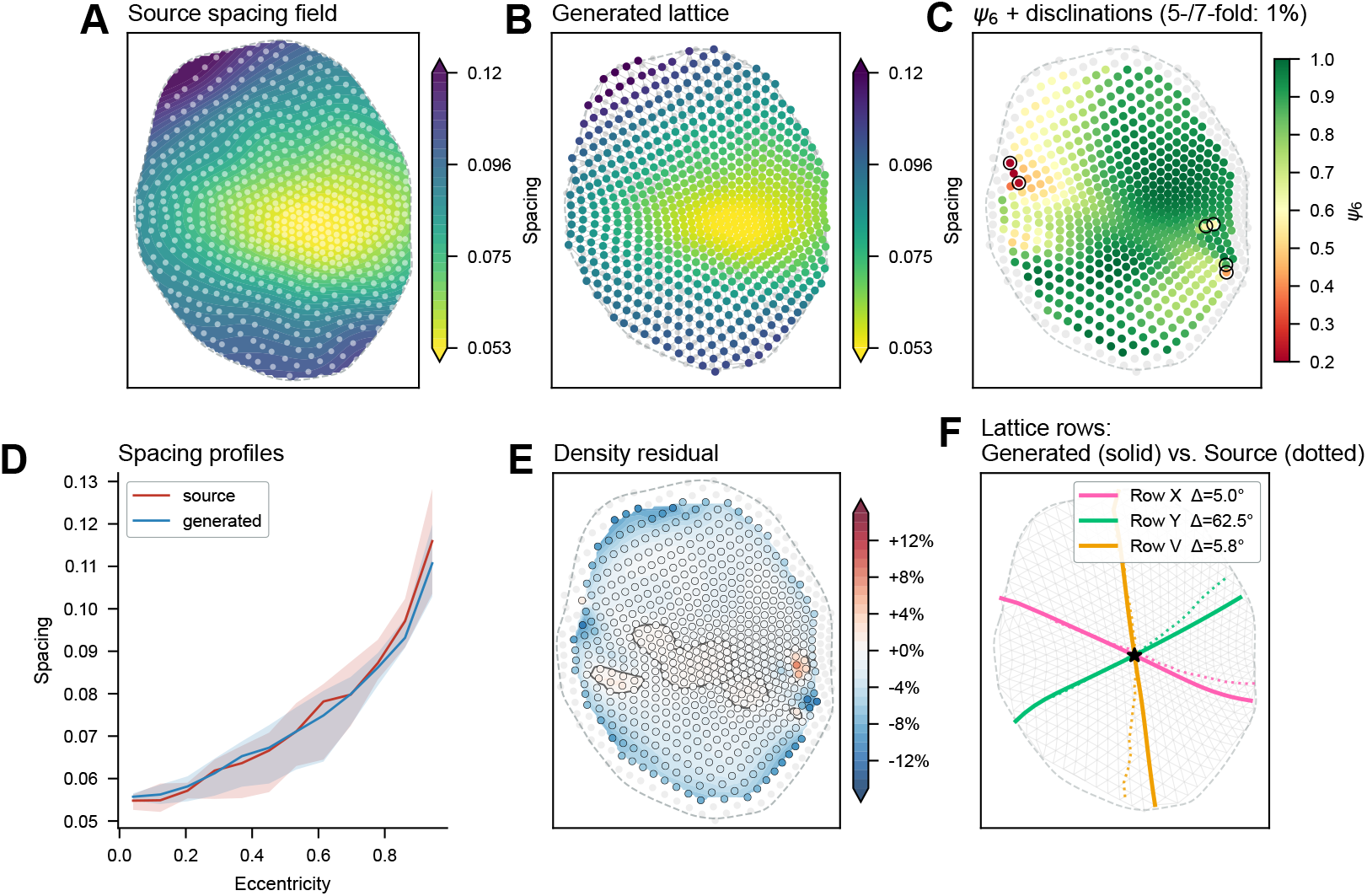
Procedural ommatidial lattice generation from empirical 2D data. **(A)** Extracted local spacing field (target ommatidial density), *N*_*target*_ = 699 ommatidia, mapped onto the 2D stereographic plane. The field is generated via Radial Basis Function (RBF) interpolation from digitised 2D source coordinates (white dots, derived from [68]). **(B)** The final generated 3D ommatidial lattice (*N*_*generated*_ = 705), colour-coded by the achieved local lens spacing. The algorithm dynamically relaxes points to match the target density while maintaining a hexagonal structure. **(C)** Hexatic order parameter (*ψ*_6_) indicating local lattice regularity. Perfectly hexagonal tiling approaches *ψ*_6_ *≈* 1 (green). Topological defects (5- and 7-fold coordinated ommatidia, i.e. disclinations) necessary to wrap the hexagonal lattice around a curved surface are outlined in black. **(D)** Radial spacing profile comparing the empirical source data (red) and the generated digital lattice (blue). Solid lines represent the median spacing as a function of eccentricity from the eye centre, with shaded regions denoting the interquartile range (IQR). **(E)** Spatial mapping of the density residual, demonstrating the minimal relative error between the generated lattice spacing and the target density field. **(F)** Preservation of lattice grain. The generated lattice rows (solid lines) accurately follow the empirical orientation traces of the source data (dotted lines) along the three primary hexatic axes.

**Figure 4:**
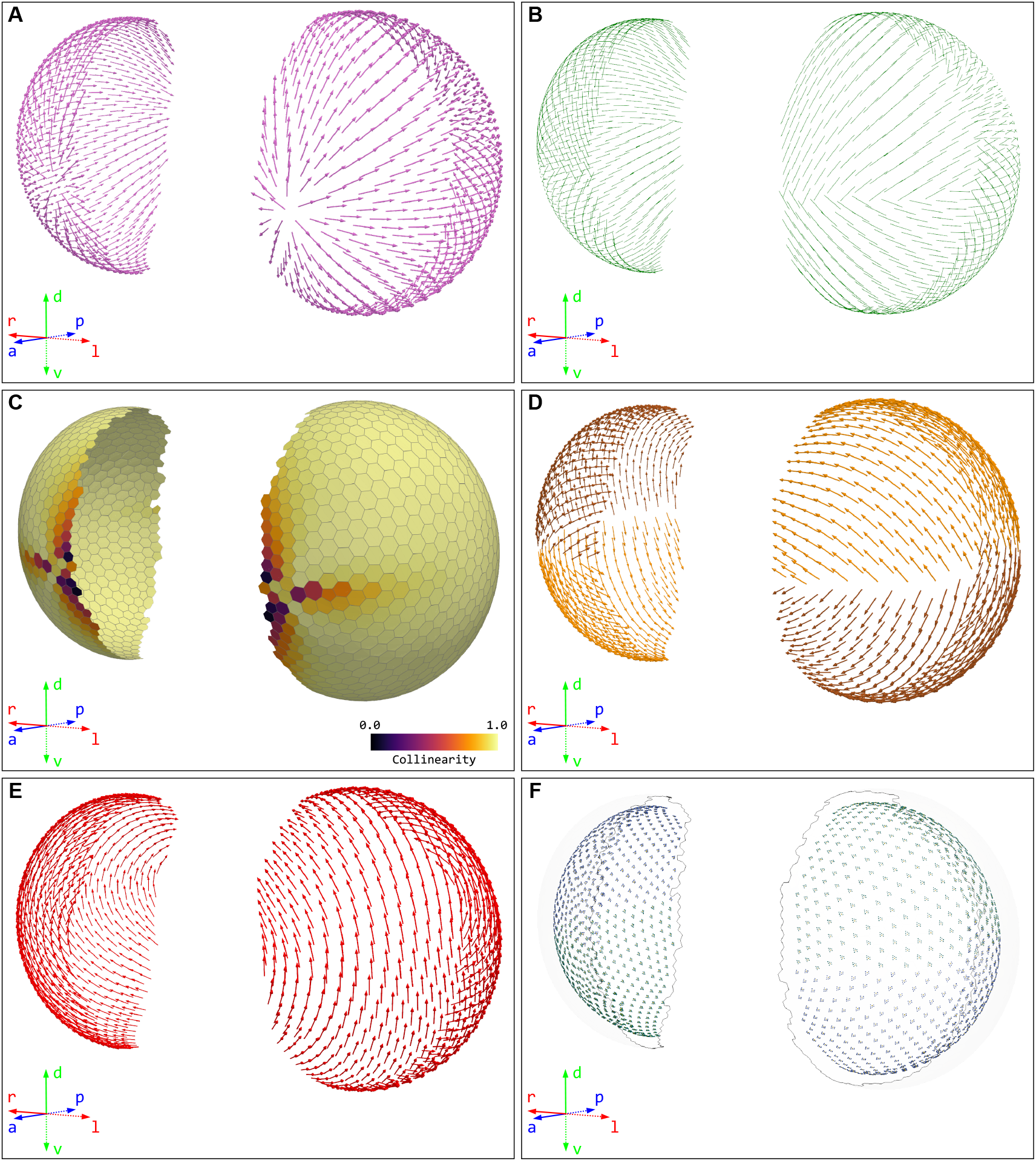
Rhabdomere bundle orientation and actuation geometry. **(A)** Projected optic flow field. World-space optic flow corresponding to forward flight (accounting for the *∼*10.1^*°*^ head pitch) projected onto the local tangent plane of each ommatidium. **(B)** Combed and smoothed alignment phasors. The reference alignment field (green lines) generated after resolving the anterior expansion singularity via diagonal combing and applying nematic *k*-NN smoothing (*k* = 8) independently within each chirality hemisphere. **(C)** Flow-alignment collinearity heatmap. Scalar projection (|**v**_flow_ *·* **v**_align_| ) quantifying local alignment between raw projected optic flow and the combed reference field. Deviations from pure flow are concentrated around the frontal pole of expansion, reproducing the frontal oblique alignment observed *in vivo*. **(D)** Structural major axes and equatorial chevron. Derived major structural axes of individual rhabdomere bundles (the R3–R6 axis in *Drosophila*), maintained at an average 81^*°*^ offset relative to the alignment field. Glyphs are colour-coded by hemispheric chirality (dorsal vs. ventral), showing the characteristic equatorial discontinuity (sharp “V-shape”) where chirality and orientation invert. **(E)** Microsaccade actuation axes. Photomechanical microsaccade actuation axes derived as an anatomical *∼* 28.6^*°*^ offset from the major structural axis (corresponding to the R1–R2–R3 axis in *Drosophila*). Global nematic smoothing and polarity alignment yield a unified, dorsally oriented actuation field across the entire eye with no equatorial discontinuity. **(F)** Rhabdomere bundles morphology. Full model showing all individual rhabdomeres. Central rhabdomeres (R7/8) are marked in gold, peripheral rhabdomeres (R1–R6) are colour-coded by hemispheric chirality (teal: unmirrored; blue: mirrored), illustrating the mirror-symmetric trapezoidal geometry across the dorsal-ventral equator.

Once this optimal yaw (*χ*) and mirror-symmetric chirality is set for every rhabdomere bundle, other ommatidial properties can be derived. First, the microsaccade actuation axes are calculated as an offset from the bundle’s structural grain. In *Drosophila*, the actuation axis is approximately 28.6^*°*^ off the bundle’s main axis [32]. Because this offset is chirality-dependent, we generate a saccade phasor field using the *±* 28.6^*°*^ rotation and perform a second nematic kNN smoothing pass. Unlike the bundle orientation, this smoothing is performed across the entire eye. The resulting axes are then polarised using the chirality information again, so that they point in a consistent sense in the head frame (i.e. dorsally for *Drosophila*). This produces a globally uniform actuation field that lacks the equatorial kink, that maintains the alignment to the structural offset (for a target angle of 28.6^*°*^, mean = 27.9^*°*^, std = 6.6^*°*^; 75 % within *±* 2.0^*°*^), matching biological observations [32]. This ensures that the simulated photomechanical movements (fast contraction and slow relaxation, see *Sub-ommatidial biophysics and dynamic optics*) are both anatomically grounded and, importantly, consistent with the underlying neural wiring (see below).

### 2.4 Cartridge wiring: simple versus neural superposition

In the simplest configuration, each ommatidium maps to a single cartridge: when *R* = 1 (a fused rhabdom, or a single central rhabdomere such as R7/8), the cartridge response is just that ommatidium’s rhabdomere response, and the combination matrix introduced earlier is the identity. Multi-rhabdomere eyes that do not pool across ommatidia can be wired the same way, one cartridge per ommatidium. The fly eye, however, is a case of neural superposition: rhabdomeres from neighbouring ommatidia that share a visual axis are pooled into a common cartridge.

With the orientation (*χ*) and chirality of every ommatidium’s bundle of rhabdomeres established, the framework must determine the neural connectivity of the cartridges (the functional units of the Lamina where R1–R6 signals from neighbouring ommatidia converge). In an ideal hexagonal lattice, this wiring follows a fixed topological rule. However, in biological eyes and procedurally generated models, local lattice irregularities, curvature, and the equatorial chirality flip make a simple index-based wiring table insufficient. A seemingly intuitive approach to wiring these cartridges would be to use a purely directional search: identifying rhabdomeres whose optical axes are most nearly parallel or converge on the same point in world space. However, such a “naive” angular overlap method is insufficient, particularly on imperfect lattices: this would allow incorrect connections to an ommatidium that is angularly close but topologically distant. In a real ommatidial lattice, neural superposition is a structural arrangement where axonal projections are constrained by the local hexagonal topography [74, 76]. To address this, we implement a two-stage spatial template-snapping and global optimisation algorithm that enforces the strict topological grain of the eye while gracefully handling lattice defects, the equator (if needed), and local distortions.

In a first stage, the algorithm generates a set of plausible wiring candidates for each “home” ommatidium. We generate a 2D projection of the ideal rhabdomere bundle into the ommatidium’s tangent plane, rotated by the local rhabdomere bundle yaw (*χ*_*i*_) and mirrored according to the local chirality. To account for local curvature and lattice variations, neighbour distances are normalised by the spacing of the local first-ring neighbours, effectively mapping the search into dimensionless “lattice units”. The algorithm then performs a bounded search for a similarity transform *w* = *s · e*^*iθ*^ (representing minor deviations in scale and rotation) that aligns the ideal template with valid surrounding ommatidia. For each tested transform, a Hungarian algorithm (linear sum assignment, [82]) resolves the optimal one-to-one mapping between the peripheral rhabdomere slots and the neighbouring ommatidia. Any assignment exceeding a tight distance threshold (e.g. 0.5 lattice units) is rejected. If a highly distorted region yields no valid rigid transforms, an adaptive fallback allows a wider search radius without a rigid transform. The top-scoring candidates (that correctly “snap” to the local neighbourhood) for each lens are saved for the next stage.

In the second stage, the greedy, per-lens assignments are unified into a globally consistent wiring using a Mixed-Integer Linear Programming (MILP) solver. This global optimisation fulfils a critical requirement: a given rhabdomere projects to a single cartridge, and conversely, each cartridge only takes one rhabdomere per neighbouring ommatidium. The solver minimises a cost function balancing the total spatial assignment error (Euclidean distance of the snaps) against penalties for heavily distorted orientations or dropped connections. This global optimisation produces a coherent neural superposition pathway that strictly adheres to local topography, resolves overlapping claims competitively, allows partial cartridge wiring at the edges of the eyes and along the equatorial chirality inversion, and is robust to any structural noise such as procedural defects, missing lenses, etc.

The two-stage MILP solver fixes *which* neighbouring lenses feed each cartridge, but the bundle yaw (*χ*_*i*_) used to generate the wiring candidates was itself derived from the smoothed optic-flow alignment field, and need not be perfectly consistent with the discrete assignment the solver ultimately chose. RhabdoForge therefore exposes an optional refinement pass that reconciles the two.

For every home lens, the optical axes of its wired donor ommatidia are projected into the home focal plane and compared, slot by slot, against the ideal template positions implied by the current *χ*_*i*_. The resulting per-ommatidium angular residual (a template-radius-weighted circular mean over the wired donors) is calculated. The algorithm can also compute a radial scale ratio or fit a continuous 2 *×* 2 anisotropic stretch matrix to account for local lattice shear. This geometric adjustment is accumulated, lightly smoothed over the first-ring lattice graph, under-relaxed, and clamped to a small maximum nudge to prevent orientation flips. Only ommatidia with a sufficient number of wired donors (e.g. *≥* 3) are trusted, the rest inherit the smoothed correction from their neighbours. The final nudge is applied to *χ*_*i*_ (and the bundle’s structural footprint, if scaling or anisotropy is enabled), after which all dependent quantities (the rotated rhabdomere offsets *δ***p**_*j*_, the actuated viewing directions, and the microsaccade axes **s**_*i*_) are rotated and recomputed to stay consistent.

Performing the discrete MILP wiring first and this geometric fit second effectively reverses the chronological sequence of biological eye development. *In vivo*, the physical lattice geometry and planar cell polarity (PCP) establish the precise rhabdomere orientations, which subsequently guide axon growth and cartridge assembly. In this procedural digital case, however, this retroactive adjustment is advantageous: because the geometric correction is driven by the specific donors the solver has already successfully linked, it neatly realigns each bundle to the true local lattice grain, and reduces the spurious orientation “zones” that a purely geometric global smoothing tends to nucleate around lattice irregularities and disclinations.

### 2.5 Sub-ommatidial biophysics and dynamic optics

Building on the per-rhabdomere acceptance model (see here), the luminance trackers (see here), and the per-ommatidium actuation axes (see here), RhabdoForge implements a sub-ommatidial biophysical cascade running in GPU compute shaders. This cascade couples the photomechanical actuation of the rhabdomeres to real-time updates of their optical properties, geometric clipping, and physical gain, and operates at every rendered time step, with a temporal resolution that can be manually defined. It consists of several non-linear stages.

#### Photomechanical microsaccades

The engine implements a biophysical reflex arc based on intracellular and optical measurements in *Drosophila* [31, 32]. The difference between the *fast* and *slow* luminance trackers (defined above) generates a local (per-ommatidium) contrast signal:

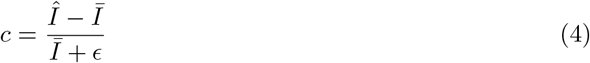

where ϵ is a small constant. Only positive contrast (luminance increments) is considered, and the reflex fires when *c* exceeds a noise threshold. The reflex is all-or-nothing: the contrast determines when a microsaccade is triggered but not how large it is, and every trigger drives the rhabdomere to full amplitude.

Kinematically, a trigger launches a committed ballistic reflex arc [31]: a fixed onset latency (*∼* 8 ms) precedes a 100 ms contraction reaching full amplitude, followed by an exponential return (*τ ≈* 190 ms) that a new trigger can interrupt, restarting the arc from the current displacement. The return time constant is the measured recovery of the rhabdomere movement (mean *∼*190 ms, [31]). The reflex sets two displacement amplitudes:

1. **Lateral displacement**: Rhabdomere tips move sideways in the focal plane along the local saccade actuation axis (**s**_*i*_), shifting the viewing direction. In *Drosophila*, lateral displacement varies from fly to fly between about 0.5 µm and 2.1 µm [32]. We use a value of 1.25 µm which sits towards the middle this range.
2. **Axial displacement**: Rhabdomeres contract longitudinally along the optical axis by 2.0 µm [32], pulling the their tips away from the lens, and closer to the focal plane.

#### Dynamic acceptance optics

In vivo, the effective acceptance angle is dynamically modulated by three distinct physical and geometric factors: axial waveguide defocus, intracellular pupil mode-stripping, and lateral aperture clipping. In RhabdoForge, the instantaneous acceptance angle Δ*ρ*_*eff*_ is calculated each frame as:

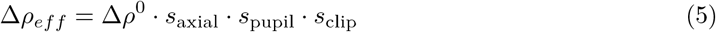

where Δ*ρ*^0^ is the dark-adapted resting acceptance angle. For the waveguide target, the sampler does not apply this product directly: the three effects are realised for each mode separately, with the pupil acting on the mode weights, the axial displacement on the mode shapes (via the pre-solved lookup), and the clipping as a scalar on the mode widths (see *The rhabdomere as the fundamental sampling unit*). The scalar ratios *s*_axial_ and *s*_pupil_ are pre-solved from the same mode solutions.

1. *Axial mode narrowing (s*_*axial*_): As rhabdomeres contract axially, their tips shift along the asymmetric defocus curve of the lens. In dark-adapted conditions, moving 2.0 µm toward focus narrows outer (R1–R6) rhabdomeres’ acceptance angles by 7 % to 10 % (*s*_axial_ *≈* 0.90–0.93) and central (R7/8) rhabdomeres’ by 16.6 % (*s*_axial_ *≈* 0.83). Crucially, because closing the pupil strips bound modes selectively, the dominant mode’s ASF changes, meaning that the axial narrowing floor depends on the pupil state. When light-adapted, R2–R5 narrow by only about 5 % and R7/8 by about 9 %, while R1 and R6 slightly widen (*s*_axial_ *≈* 1.03–1.04). This is because the side of the defocus-response minimum on which a mode falls is set by the dominant mode.
2. *Intracellular pupil mode-stripping (s*_*pupil*_): As described in *Reduction and adaptation*, the seconds-scale pupil closure attenuates higher-order modes, narrowing the resting acceptance angle to *s*_pupil_ *≈* 0.55–0.63 across all rhabdomeres.
3. *Lateral aperture clipping (s*_*clip*_): As rhabdomeres translate laterally, they displace relative to the (fixed) optical aperture and the cone/pigment-cell aperture. The cone aperture is mechanically dragged along by adherens junctions at approximately half the rhabdomere’s lateral displacement [31], which creates a vignetting effect. In RhabdoForge, this clipping is parametrised by the ratio *r*_*clip*_ *∈* [0, 1] (where 1.0 represents no clipping). The dynamic clipping factor is evaluated based on the relative offset:

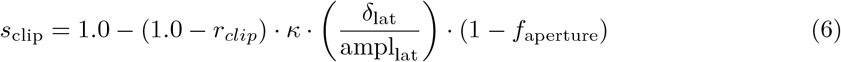

where *κ ∈* [0, 1] is the per-rhabdomere clipping weight (0 on the bundle axis, 1 for the rhabdomere farthest from the bundle centre) and *f*_aperture_ = 0.5 represents the aperture-follow fraction. Central rhabdomeres (R7/8), which are on-axis (*κ* = 0), exhibit minimal lateral excursion and are thus not subject to this lateral clipping. Note that this *r*_*clip*_ ratio is a phenomenological factor whose effect is to make the ASF narrower during lateral displacement. It can be attributed to geometric vignetting (clipping of the ASF against the lens’s diffraction footprint and the cone/pigment-cell aperture), or to any neural or voltage-level nonlinearity that narrows the effective ASF during the microsaccade. It is symmetrical rather than a true one-sided vignetting effect as observed *in vivo*, and it is calibrated rather than empirically measured. We set *r*_*clip*_ = 0.85 in the Sub-ommatidial biophysics and dynamic optics result section.

#### Photon-concentration heuristic

In addition to ASF narrowing, axial displacement concentrates optical flux within the rhabdomere waveguide [83]. The engine optionally applies a physical photon-concentration gain boost scaling with the inverse square of the axial optical narrowing:

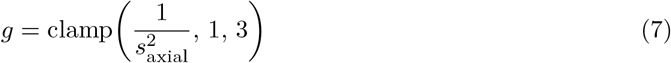

*·*

This boost is blended with unity gain via a user-controllable weight *β∈* [0, 1] (*G*_photo_ = (1 *− β*) + *β g*). Importantly, this gain boost depends strictly on the axial geometric scale *s*_axial_: it excludes pupil mode-stripping (*s*_pupil_) and lateral clipping (*s*_clip_) to avoid double-counting the optical transmittance losses handled during the GPU reduction pass (see *Reduction and adaptation*). **Retinal movement**: Finally, in addition to local rhabdomeres microsaccades, RhabdoForge supports simulating macro-scale movements of the entire retinal sheet driven by extra-ocular muscles [84]. This provides an additional layer of active-sensing control, allowing agents to execute coordinated bilateral or unilateral retinal shifts.

#### Parameter provenance

Table 3 lists the main parameters of the example *Drosophila* model used in this paper, together with their source and status, to make clear which values are taken from the literature, which are derived from the model, and which are assumed or calibrated. Note that these values document the configuration we used and not the framework: they are all user-configurable, and not imposed by RhabdoForge.

**Table 3:**
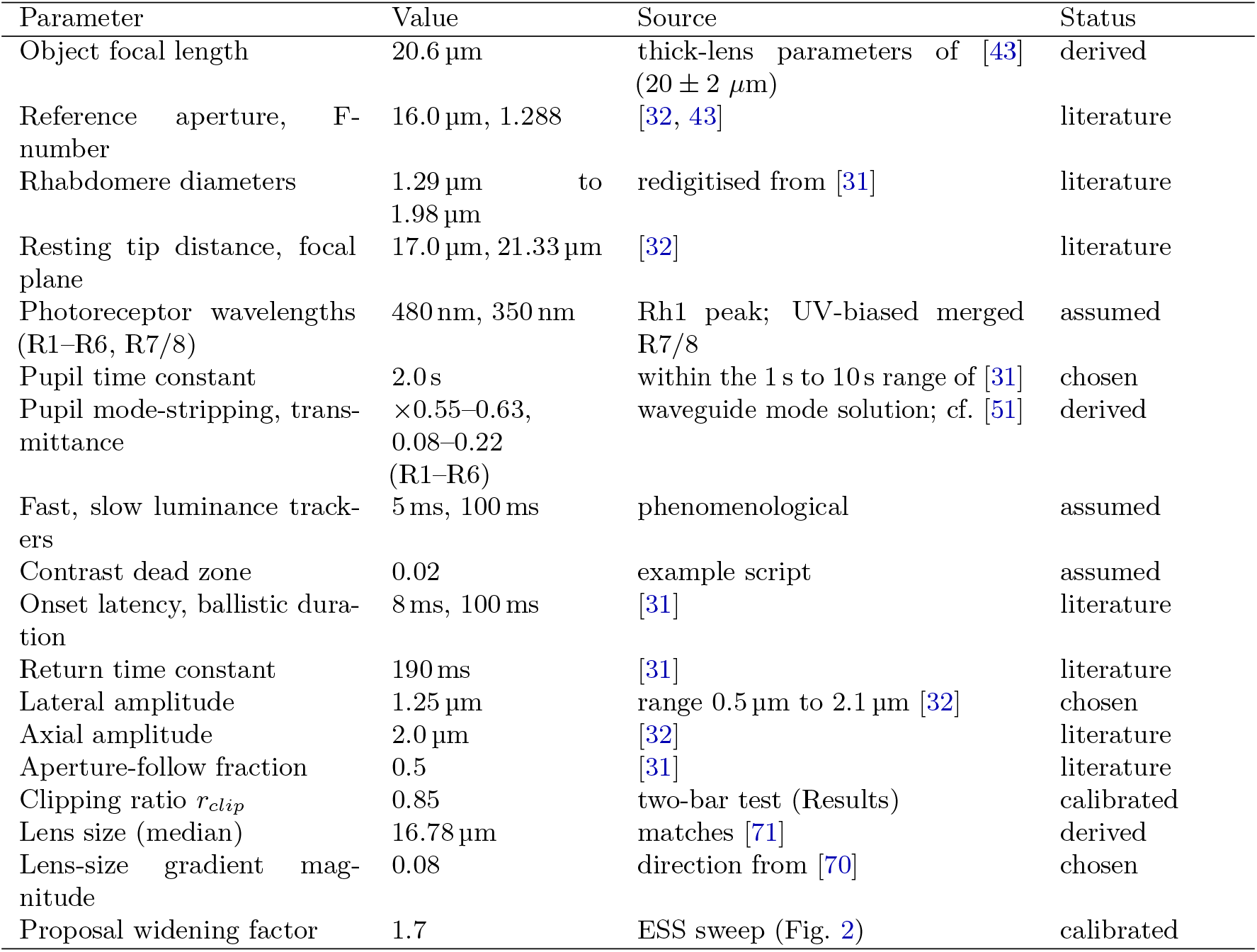
Main parameters of the example *Drosophila* model used in this paper. *Literature*: taken from the cited work; *derived*: computed by the model; *chosen*: a value within a measured range; *assumed*: no direct source; *calibrated*: set using a parameter sweep or assay.

| Parameter | Value | Source | Status |
|---|---|---|---|
| Object focal length | 20.6 µm | thick-lens parameters of [43] (20 ± 2 μm) | derived |
| Reference aperture, F-number | 16.0 µm, 1.288 | [32, 43] | literature |
| Rhabdomere diameters | 1.29 µm to 1.98 µm | redigitised from [31] | literature |
| Resting tip distance, focal plane | 17.0 µm, 21.33 µm | [32] | literature |
| Photoreceptor wavelengths (R1–R6, R7/8) | 480 nm, 350 nm | Rh1 peak; UV-biased merged R7/8 | assumed |
| Pupil time constant | 2.0 s | within the 1 s to 10 s range of [31] | chosen |
| Pupil mode-stripping, trans-mittance | ×0.55–0.63, 0.08–0.22 (R1–R6) | waveguide mode solution; cf. [51] | derived |
| Fast, slow luminance track-ers | 5 ms, 100 ms | phenomenological | assumed |
| Contrast dead zone | 0.02 | example script | assumed |
| Onset latency, ballistic dura-tion | 8 ms, 100 ms | [31] | literature |
| Return time constant | 190 ms | [31] | literature |
| Lateral amplitude | 1.25 µm | range 0.5 µm to 2.1 µm [32] | chosen |
| Axial amplitude | 2.0 µm | [32] | literature |
| Aperture-follow fraction | 0.5 | [31] | literature |
| Clipping ratio *r*<sub>*clip*</sub> | 0.85 | two-bar test (Results) | calibrated |
| Lens size (median) | 16.78 µm | matches [71] | derived |
| Lens-size gradient mag-nitude | 0.08 | direction from [70] | chosen |
| Proposal widening factor | 1.7 | ESS sweep (Fig. 2) | calibrated |

### 2.6 Agent-level encapsulation and control

The components described above (eyes, with their dynamic rhabdomeres and cartridge wiring) are encapsulated at the agent level: an agent (which owns one or more eyes) that can be moved through the world under direct interactive control, along a predefined path (which can be run offline), or under closed-loop control.

To support active sensing experiments while minimize the engineering overhead for neuroscientists, RhabdoForge abstractions mirror those of most video game engines. Kinematics are controlled via an object-oriented transformation API. This allows defining active-sensing behaviours (e.g. head saccades, translational movements) for the agents, or animating any scene element or light source, entirely natively within Python, abstracting the underlying OpenGL state management. The system supports the use of pre-defined paths via a Curve and Trajectory API (useful for example for loading harmonic radar tracking data of honeybees, manually traced ant learning walks, or full foraging routes recorded with video tracking [85–87]). Furthermore, RhabdoForge features an optional, interactive “human-in-the-loop” visualisation context: one can seamlessly toggle between the agent’s multi-viewpoint ommatidial perspective (first person) and third-person observer camera in real-time 6, with full control of the agent via keyboard and mouse (or gamepad) for human-guided navigation.

#### 2.6.1 Dual running mode: latency versus throughput

A major bottleneck in closed-loop simulations for neuromorphic modelling is the latency involved in transferring rendered frames from GPU memory (VRAM) back to host RAM. RhabdoForge addresses this via a dual running mode architecture optimised for either latency (synchronous) or throughput (asynchronous).

For strictly synchronous closed-loop simulations, where an agent’s next action depends immediately on the current visual input, the renderer uses a two-stage readback: while one Pixel Buffer Object (PBO) is being filled by the GPU, the previous frame is mapped to the CPU in a second PBO, avoiding stalling the PCIe bus.

For offline tasks, where for instance an agent follows a pre-defined trajectory, RhabdoForge can be used in asynchronous mode: frames are rendered into a circular History SSBO on the GPU without blocking the Python host. Data is only transferred back to the CPU in large batches (e.g. every 1000 frames). This significantly reduces the per-frame overhead of the PCIe bus and allows the renderer to achieve much higher performance.

### Additional implementation details

#### 2.7.1 Host-side logic and an open, hardware-agnostic GPU layer

To support dynamic closed-loop experiments, RhabdoForge manages all high-level scene logic, agent kinematics, and the construction of the sensory arrays directly within the Python environment on the host CPU, while all graphical and massively parallel computations (ray-scene intersections) are offloaded to GLSL compute shaders on the GPU.

We provide a custom GLSL loader, extended to support #include directives, which makes the GPU code more modular and easily extensible by researchers who want for instance to write their own custom light transport or neuromorphic filters directly on the GPU. By using an open-source and hardware-agnostic bounding volume hierarchy (BVH) library [40] and standard OpenGL compute shaders instead of proprietary ray-tracing frameworks, we avoid vendor lock-in and ensure the software is fully deployable across diverse hardware systems (being high-end workstations or standard consumer laptops).

#### 2.7.2 Rendering modes

RhabdoForge supports two distinct rendering modes to balance speed and biological realism. The ray-tracing mode is the primary one. It is optimised for real-time, closed-loop navigation. It performs a single-hit intersection test for each ray, returning the surface colour and texture information. This mode is highly efficient, capable of rendering point cloud environments containing *∼* 10^7^ primitives (LiDAR datasets from Habitat3D [88]) at hundreds of frames per second. Just like other existing compound eye simulators [34, 35], this single-hit ray-casting mode efficiently determines primary visibility but omits secondary light transport.

For studies in visual ecology where ambient occlusion, soft shadows, and other types of complex light transports are required, RhabdoForge also includes a stochastic multiple-bounce path-tracing mode [46, 47]. This mode uses Monte Carlo sampling to simulate light bouncing within the environment. While more computationally expensive, by evaluating recursive ray bounces, the engine naturally resolves global illumination, ambient occlusion, colour bleeding between nearby surfaces and other phenomena, subjecting the virtual insect to an environment that is closer to the complex lighting characteristics of natural habitats.

#### 2.7.3 CPU-side spatial queries

As detailed above, RhabdoForge relies on a Bounding Volume Hierarchy (BVH) spatial acceleration structure for 3D rendering. But by maintaining a persistent reference to the C++ BVH objects within the host system’s memory, and in addition to BVH’s role in the GPU rendering, CPU-side intersection tests and spatial queries can also be executed directly from Python. This enables low-latency physical interactions, including proximity sensing and basic collision detection (via ray- or sphere-casting), without external dependencies. By reusing the same acceleration structure for GPU visual rendering and CPU spatial queries, the framework avoids the computational overhead of running a separate physics engine in parallel.

## 3. Results

### 3.1 Closed-loop validation: Optic flow-based centring response

#### 3.1.1 Experimental paradigm

As a small proof of concept, we replicated the classic “tunnel centring response”, observed in honeybees (*Apis mellifera*) navigating narrow corridors [89] that balance the bilateral optic flow to fly in the centre of the tunnel. This is a simple, yet good experiment to demonstrate the ability of RhabdoForge to maintain high-frequency interaction between a biological sensory periphery and a downstream neural controller in closed-loop neuroethological simulations.

In this experiment, an agent was equipped with a honeybee eye model (using an adapted version of Stürzl’s model [67]), a single output value per ommatidium (fused rhabdomeres), and no active microsaccades. Each of the two eyes consisted of 3846 ommatidia. The agent was placed in a long virtual tunnel (0.2 *×* 0.2m cross-section) textured with high-contrast, random irregular chequer patterns. To test the robustness of the centring response, we performed multiple trials with randomised starting positions across the entrance plane (*x* and *y* coordinates).

A simple proportional controller balanced the mean Elementary Motion Detection (EMD) response (see below) from the left and right eyes to modulate either the agent’s yaw (steering), or lateral position (strafing). The lateral EMDs were configured to be sensitive primarily to horizontal flow.

#### 3.1.2 Elementary motion detection model

The first agent’s “brain” was implemented with a bilateral array of EMDs, based on the standard Hassenstein-Reichardt Correlator (HRC) [90, 91], to estimate local optic flow. The motion detection pipeline was partitioned between the GPU-side preprocessing and Python-side neural logic. The GPU-side consisted of the temporal accumulation (low-pass filter) as described in the *Reduction and adaptation* section. On the Python side, for each neighbouring ommatidial pair (*i, j*) (because *R* = 1 in this model), the motion response *R* was calculated as the high-pass filtered luminance signal *L* cross-correlated with a delayed version of its neighbour.

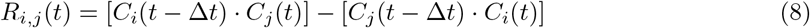

where *C* represents the contrast-adjusted signal. This signal is derived from the raw ommatidium’s luminance *L* via a high-pass filter (approximating the luminance adaptation observed in the insect lamina, [92]): 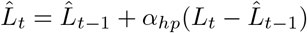, with 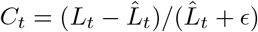 . The temporal delay Δ*t* was implemented as a first-order low-pass filter with time constant *τ*_*delay*_. While this model is a simplified abstraction of the optic lobe, it serves as a robust benchmark for evaluating real-time sensory-motor loops.

#### 3.1.3 Gradient-based optic flow model

To address the spatial frequency dependence of the HRC model (see below), a second motion detection pipeline was implemented using a gradient-based optic flow estimator [93]. This model calculates true angular velocity via the local optic-flow constraint, effectively cancelling out contrast and spatial frequency [94]. For each ommatidium (again, because here *R* = 1), the temporal gradient *I*_*t*_ = (*I*(*t*) *− I*(*t −* Δ*t*))*/*Δ*t* and the spatial gradient *I*_*x*_ = *I*_*target*_(*t*) *− I*(*t*) along the horizontal axis were extracted directly from the luminance signal *I*. To compute a robust, single-eye velocity estimate, the model aggregates these local gradients using a pooled Lucas-Kanade method:

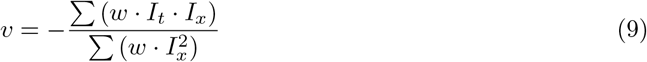

where *w* represents the projection weights of the directed ommatidial neighbours. Because this ratio-based formulation balances the temporal and spatial derivatives, it provides a direct estimate of image speed that is theoretically independent of wall texture density, offering an alternative mechanism for mimicking the texture-invariant speed estimation of real bees.

#### 3.1.4 Behavioural output

Across all trials and both control regimes, the agents converged onto the tunnel midline from randomised entry positions (Fig. 7). Centring emerged purely from balancing the bilateral mean EMD response (Fig. 7): as the agent drifted toward one wall, the rising angular velocity on that eye increased its correlator output and the proportional controller restored balance. The response was robust to additive motor noise and largely independent of control mechanism; non-holonomic (yaw) or holonomic (strafe) controllers, differing only in approach dynamics.

**Figure 5:**
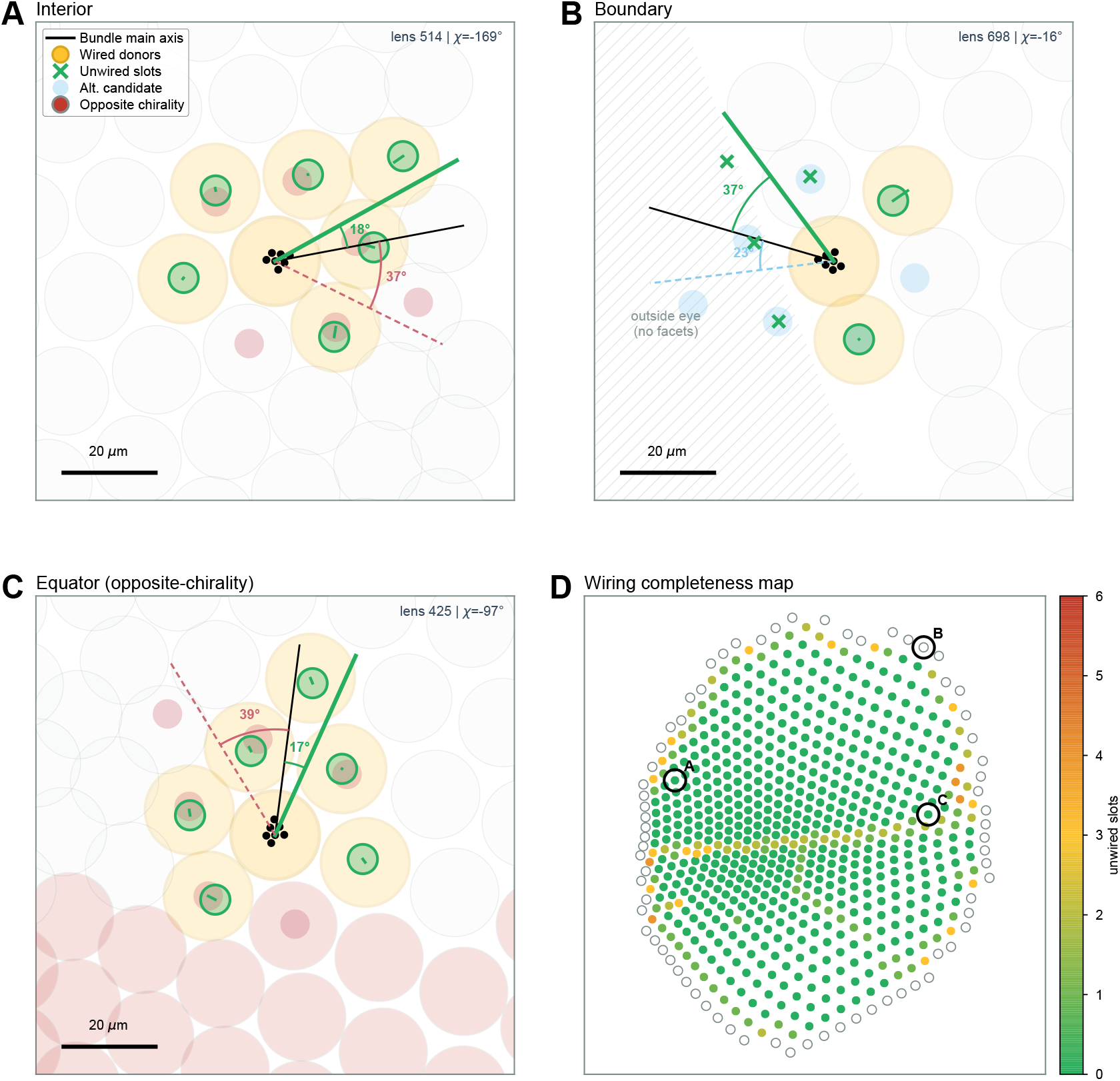
Neural superposition wiring algorithm and topological robustness. (A-C) Per-cartridge wiring diagnostics resolving the connectivity between a central ‘home’ ommatidium and its neighbours. The home ommatidium’s rhabdomeres are displayed as black dots. The solver evaluates multiple orientation-adjusted candidate templates (coloured lines/arcs) to find the rigid similarity transform that best snaps peripheral rhabdomere slots to the local lattice. Successfully wired donor ommatidia are highlighted in yellow. Unwired slots on the winning template are marked with a green cross. Alternative, rejected candidate templates are shown in translucent colours. **(A)** An interior region displaying a complete, rigid structural match. **(B)** The boundary of the eye, where the algorithm gracefully handles the edge of the optical field by leaving corresponding cartridge slots unwired rather than forcing anatomically incorrect spatial distortions. **(C)** The equator, illustrating the algorithm handling the abrupt structural chirality inversion. Immediate neighbours with opposite chirality are outlined in red and excluded from the local neural superposition wiring. **(D)** Global retinotopic wiring-completeness map of the modelled *Drosophila* eye. Colours denote the number of unwired (dropped) slots per cartridge. While the eye boundary naturally loses connections, the interior and the equatorial discontinuity maintain high completeness despite localized topological defects. Black circles indicate the locations of the exemplar lenses detailed in A–C.

**Figure 6:**
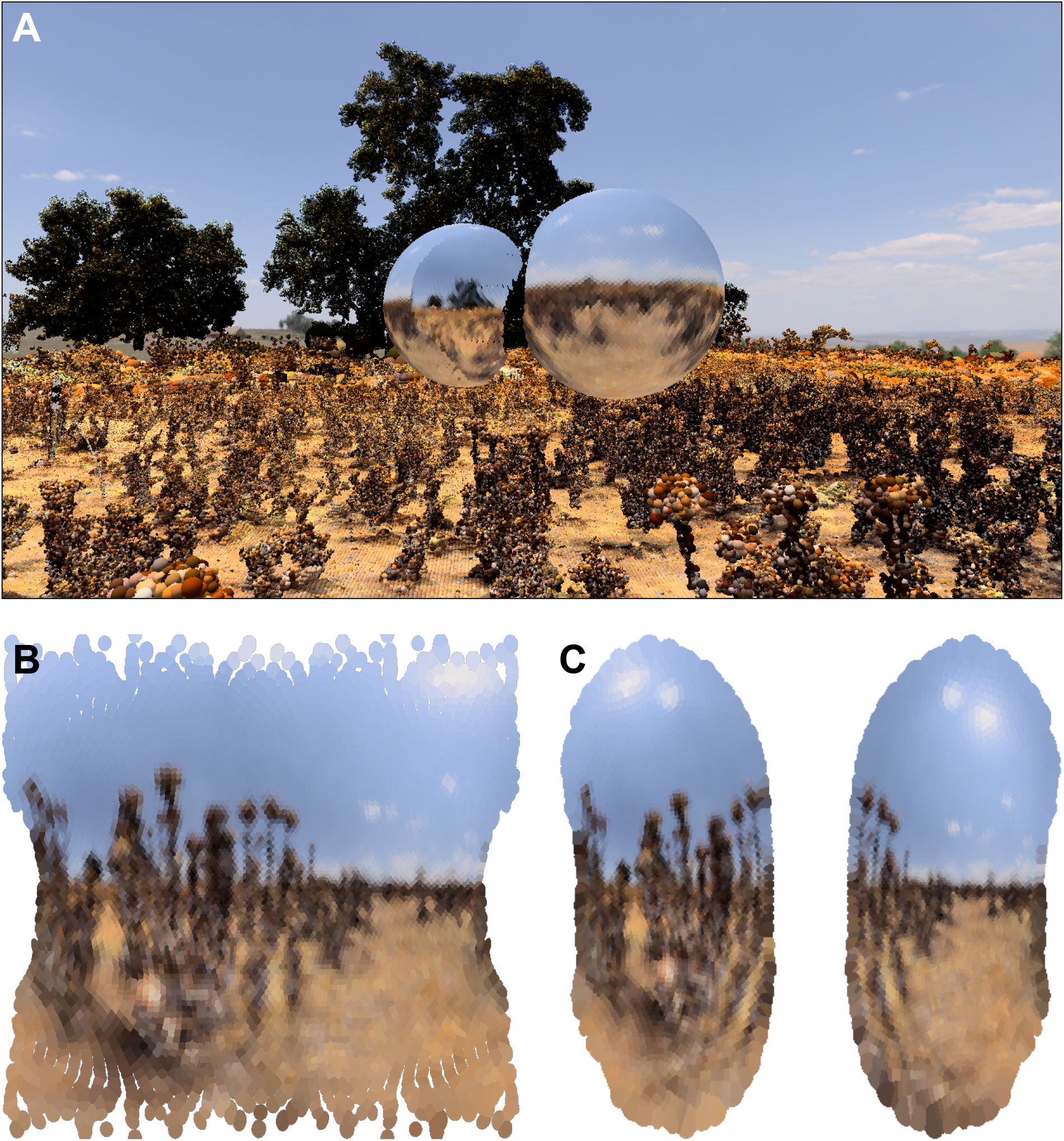
Observer perspective and first-person visual projections in complex 3D environments. Simulation of a honeybee (*Apis mellifera*) visual system (*N* = 7,692 ommatidia) navigating the *Seville* terrestrial LiDAR dataset from Habitat3D (*∼* 10^7^ primitives, [88]). Note that the honeybee possesses fused rhabdoms, whose photoreceptors share a central optical axis: RhabdoForge returns one spectral response per photoreceptor, but for display here these are pooled into a single integrated colour response per ommatidium. **(A)** Third-person observer perspective. External view showing the compound eyes in the 3D environment. Each ommatidium is individually ray-traced and shaded by its integrated radiance, reflecting the local scene geometry, ground vegetation, and sky. **(B)** Anatomical position projection. First-person visual output mapped onto the 3D surface coordinates of the left and right compound eyes. This view preserves the physical separation, head-space morphology, and ocular boundaries of the eyes. **(C)** Retinotopic visual-field projection. Radiances unprojected along individual viewing directions onto spherical coordinates. Each ommatidium is drawn with its ASF footprint, illustrating the resulting panoramic visual field, local sampling density variations, and frontal binocular overlap.

**Figure 7:**
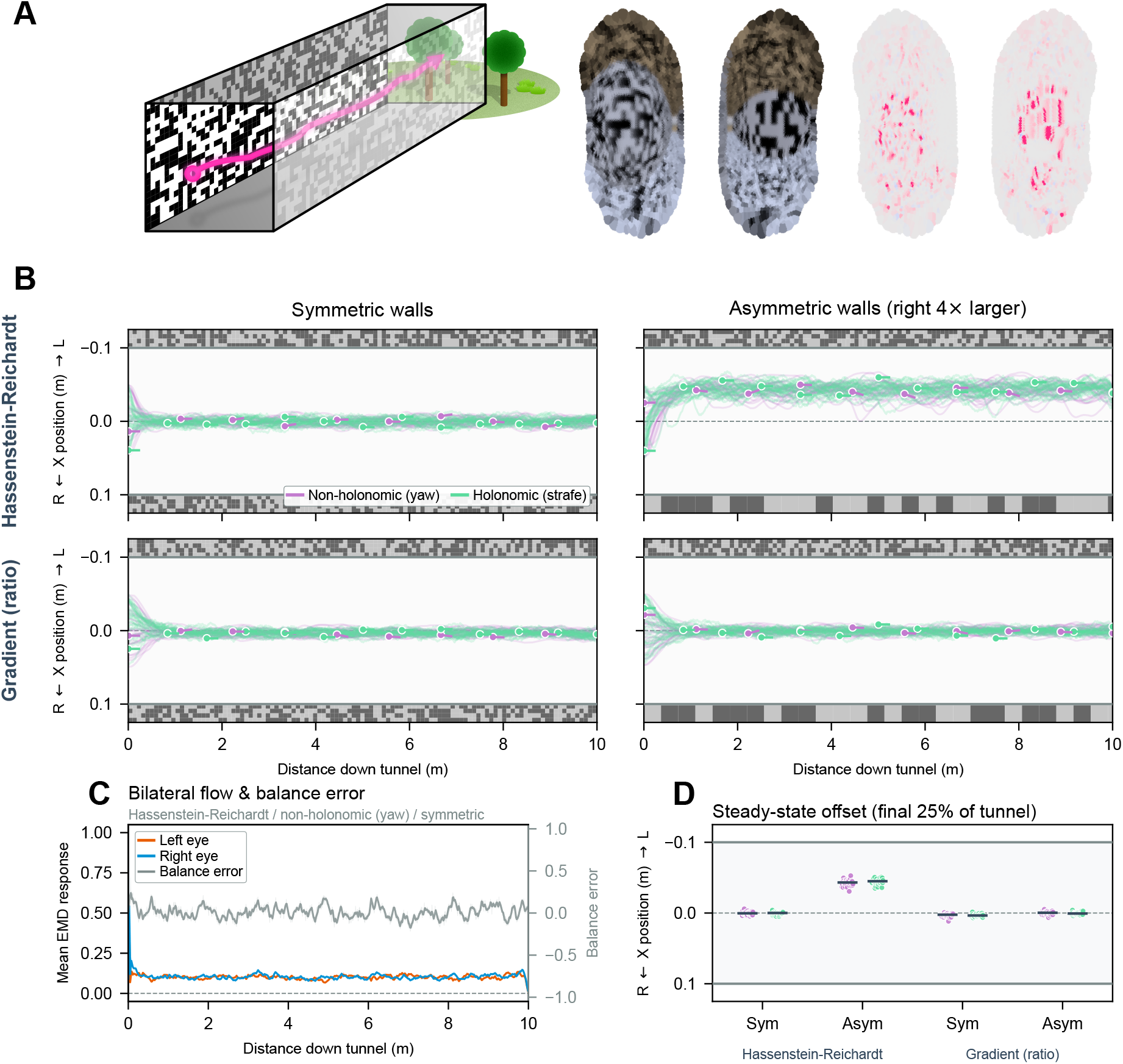
Real-time closed-loop active sensing: Optic flow-based tunnel centring. **(A)** Left, Schematic representation of the virtual tunnel environment.; Right, the simulated bee’s first-person view in colour and as a heatmap of optic flow (magenta, intensity intensity *∝* flow (*a*.*u*.). In these views, walls are asymmetrical in density. **(B)** Overhead view of flight trajectories (n = 30 per condition) starting from randomised lateral entry points (on the left of schema). Rows compare environments with symmetric vs. asymmetric wall textures (a 4-fold difference in chequer spatial density). Columns compare visual motion processing models: the contrast-dependent Hassenstein-Reichardt Correlator (HRC) and a gradient-based flow detector that estimates true angular velocity. Colours distinguish the control scheme used by the agent (purple: non-holonomic yaw steering; green: holonomic strafe). Markers denote sampled agent headings. **(C)** Smoothed bilateral motion responses and subsequent balance error over time for a representative HRC trial in the symmetric tunnel. **(D)** Steady-state lateral offset, measured over the final 25 % of the tunnel length for all trials (dots represent individual trials; horizontal lines denote the median). While both models centre correctly in symmetric tunnels, only the gradient-based model maintains spatial-frequency invariance in asymmetric conditions, successfully recovering the texture-independent centring behaviour of real honeybees [89]

We then probed the response with an asymmetric condition in which the two walls carried random chequer patterns differing four-fold in density. When relying on the HRC model, the agent settled off-axis, closer to the higher-spatial-frequency (denser) wall, in clear contrast to real bees, whose centring is invariant to such four-fold differences, and is displaced only when a grating is physically set in motion [89]. This divergence is a direct consequence of the motion model: forward translation past a pattern of period *P* produces a contrast frequency *V/P* that is independent of viewing distance, and the two walls present intrinsically different contrast frequencies, fixed by their period ratio. Unable to balance the two signals at the midline, the HRC agent instead settles where the distance-dependent component of the correlator response compensates for the spatial-frequency difference, i.e. closer to the higher-frequency side. This fits with the Hassenstein-Reichardt correlator’s known dependence on contrast frequency rather than true angular velocity [95, 96]. In contrast, when the agent was controlled by the gradient-based optic flow model, it successfully maintained a centred trajectory along the midline, invariant to the four-fold difference in wall texture density. Because the gradient model estimates true angular velocity by evaluating the ratio of temporal and spatial derivatives, the intrinsic spatial-frequency difference between the walls cancels out. This allowed the agent to balance the bilateral signals at the geometric centre of the tunnel, recovering the texture-independent centring behaviour of real bees [89].

#### 3.1.5 Computational performance

The entire perception-action loop (comprising GPU-accelerated ray-tracing of 7,692 ommatidia with 256 samples each, data transfer, and Python-based EMD evaluation) was executed comfortably in real time at a fixed simulation rate of 100 Hz. On standard consumer-grade GPU hardware, the combined rendering and evaluation latency accounted for only a small fraction of the allocated 10.0 ms inter-frame window. This low computational overhead confirms that RhabdoForge leaves sufficient GPU and CPU headroom to support more complex downstream neuromorphic architectures (e.g. Spiking Neural Networks) without compromising real-time, closed-loop interaction.

### 3.2 Sub-ommatidial biophysics and dynamic optics

Recent studies demonstrate that insects can resolve spatial details finer than their interommatidial angle, a phenomenon that the classic view of the compound eye as a static array of “pixels” fails to capture [31, 32]. This hyperacuity phenomenon is thought to be achieved through photomechanical microsaccades: rapid, luminance-driven photomechanical twitches of the rhabdomeres. The effect of microsaccades also interacts dynamically with the one of the intracellular pupil and the fixed optical aperture. To demonstrate RhabdoForge’s capacity to simulate this entire cascade, we implemented a two-bar hyperacuity-style assay and used it to decompose the dynamics into their interacting constituents.

#### 3.2.1 Experimental paradigm

Because *Drosophila* photomechanical microsaccades move the rhabdomeres along the local R1–R2–R3 axis [32], which is approximately vertical in visual space in a forward-pointing cartridge, we used a stimulus of thin, high-contrast horizontal bars (*±*1.0^*°*^ wide) presented on a frontal plane 2.0 m ahead of the agent, and oscillated the agent vertically ( 1.0 m linear sweep, *≈±*40 ^*°*^*·* s^*−*1^ apparent angular speed at the centre of the field) to induce apparent motion orthogonal to the bars. We contrasted a single-bar control with a two-bar stimulus separated by 4.0^*°*^ centre-to-centre (a 3.0^*°*^ gap). The virtual eyes used the complete *Drosophila R* = 7 rhabdomere bundle (R1–R6 plus the fused central R7/8), with neural-superposition wiring enabled, and the readout was taken from a single forward-pointing cartridge (lens aperture 17.7 µm, with a resting R1–R6 Δ*ρ* of 4.56^*°*^ to 5.19^*°*^). The biophysical cascade was run at a 1.0 ms biological time step. To isolate the effect of ASF geometry, the physical photon-concentration gain was disabled (*β* = 0) so that any change in resolving power arises from the ASF dynamics alone. Each rhabdomere cast 512 rays per frame.

To dissect the interacting mechanisms, the forward cartridge was recorded across several states. First, we forced the intracellular pupil into three steady-state adaptation levels (*dark-adapted, halfway*, and *light-adapted*), which progressively narrow the baseline ASF by stripping higher-order waveguide modes. Second, for each pupil state, we decomposed the microsaccade into four kinematic conditions: *none* (static ASF, no movement at all), *axial* (axial displacement only, narrowing the ASF via waveguide defocus with no change in viewing direction), *lateral* (lateral displacement only, translating the viewing direction while triggering the aperture clipping), and *full* (the complete microsaccade). We recorded the simulated radiance reaching the rhabdomeres of the target cartridge across the sweep, for each individual peripheral rhabdomere type (R1–R6), and for their pooled signal.

#### 3.2.2 Spatiotemporal encoding and dynamic ASF narrowing

To dissect how these dynamics influence resolving power, we consider the two geometric components of the microsaccade separately (see *Sub-ommatidial biophysics and dynamic optics*). The *axial* component moves the rhabdomere tip along the optical axis, and the waveguide solver evaluates the resulting acceptance angle at the displaced defocus. The *lateral* component shifts the viewing direction by 1.25 µm and clips the ASF against the lens aperture and the cone/pigment-cell aperture, which follows the rhabdomere at about half its lateral displacement [31, 32]. This clipping phenomenon is represented in RhabdoForge by the phenomenological clipping ratio *r*_*clip*_ (the ASF scale at full relative lateral offset, with *r*_*clip*_ = 1.0 for no clipping). In vivo, this clipping is likely asymmetric while our ratio is symmetric, but the qualitative effect is what matters here. We set *r*_*clip*_ = 0.85 using this same two-bar test, as a value giving a moderate but detectable effect in the dark-adapted state.

Firstly, the vertical sweep produces a direction-dependent offset between the response profiles for upward and downward stimulus motion (Fig. 8D). This offset is carried by the lateral component of the microsaccade, which displaces the viewing direction along the sweep axis: a stimulus met head-on by the saccade produces a compressed, shorter-latency response compared to one being chased by the saccade. Therefore, the responses for the two stimulus directions are pushed in opposite directions (about 0.20^*°*^) for the *lateral* and *full* conditions, while the purely *axial* component produces none (about 0.03^*°*^). This asymmetric readout shows that the per-rhabdomere lateral dynamics are present, correctly oriented, and running at saccadic speeds.

**Figure 8:**
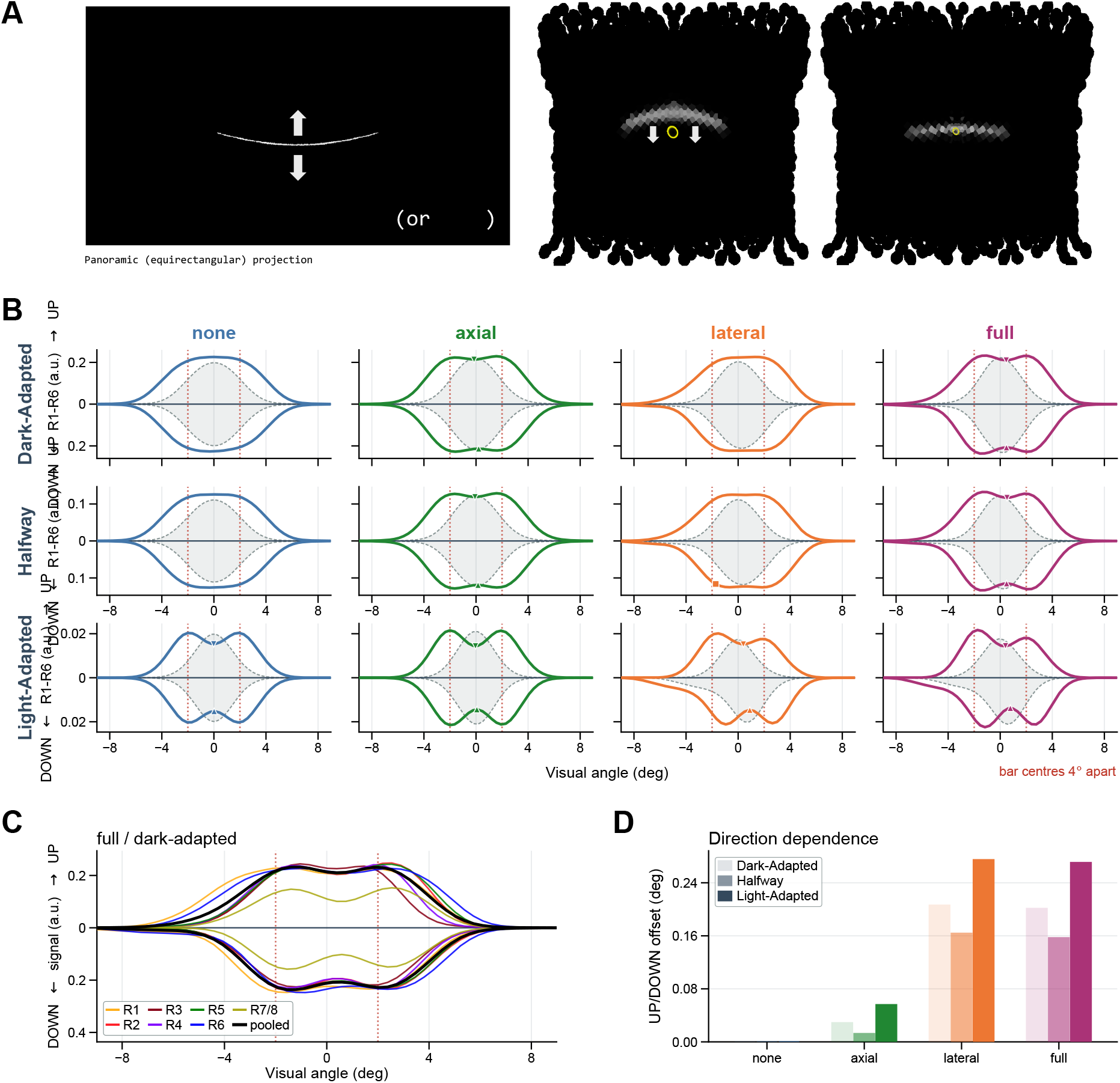
Sub-ommatidial biophysics: Pupil adaptation and microsaccadic hyperacuity. **(A)** Experimental setup. The agent sweeps vertically past high-contrast horizontal bars (1.0^*°*^ wide, 4^*°*^ separation) while the visual response from a single forward-pointing cartridge is recorded. Left: Schematic of the test, showing the single-bar condition as a panoramic (equirectangular) projection, with the two-bar condition as inset. Arrows denote the vertical apparent motion of the bars. Right: First-person view of the stimulus at two successive time steps, bar going down (arrows), with the recorded cartridge’s ASF highlighted in yellow. When the bar is detected by this cartridge, its ASF narrows. **(B)** Time-collapsed, pooled R1–R6 spatial response profiles. The upper halves (positive y-axis) show the temporal response during the UP sweep trajectory; the lower halves (negative y-axis) show the DOWN sweep. Dashed outlines denote a single-bar control stimulus; solid coloured lines represent the two-bar stimulus. Triangle markers indicate where the model successfully resolves the two bars. Rows show the intracellular pupil adaptation state (mode-stripping); columns isolate the microsaccade kinematic components. The purely *axial* component narrows the ASF to open a shallow dip; the *lateral* component alone does not resolve the bars. When combined in the *full* condition with geometric aperture clipping (*r*_*clip*_ = 0.85), spatial resolution is enhanced beyond the axial component alone. Pupil closure further deepens the dip. **(C)** Per-rhabdomere temporal responses (R1–R6 and R7/8) underlying the pooled cartridge response (black line) for the exemplar dark-adapted / *full* microsaccade condition. **(D)** Direction-dependent temporal encoding. The lateral displacement translates the viewing direction along the sweep axis, pushing the centre-of-mass of the response profiles in opposite directions (*∼* 0.20^*°*^ offset) depending on stimulus motion (UP vs. DOWN). The static (*none*) and purely *axial* conditions produce no temporal offset.

##### Axial narrowing alone partially resolves spatially

In the selected cartridge, with the anatomical 2.0 µm axial displacement, and with no extra clipping (*r*_*clip*_ = 1.0), the R1–R6 acceptance angle Δ*ρ* shrinks to about *×*0.90 its rest value (from 4.56^*°*^ to 5.19^*°*^ at rest to 4.02^*°*^ to 4.75^*°*^). This opens a shallow but detectable dip in the response for the two bars (Fig. 8B).

##### Lateral motion smears the profile unless clipped

Without clipping, the response with lateral displacement alone remains fused, and combining it with the axial displacement (*full*) is even weaker than axial alone: the lateral displacement smears the spatial profile. The direction-dependent signature of the two-bar stimulus is itself distinct from the one-bar signature, even where the time-collapsed spatial profiles are not separable. In other words, while lateral motion without sufficient clipping degrades *spatial* resolvability, the microsaccade still encodes the two-bar configuration as a temporal/direction-dependent code, consistent with the microsaccadic-sampling view in which motion converts spatial detail into resolvable temporal structure [31]. To recover static spatial resolution from the lateral displacement, clipping is required. Lateral displacement alone starts to resolve the bars spatially with *r*_*clip*_*≈* 0.7. With the chosen *r*_*clip*_ = 0.85, the lateral displacement thus still shows no dip, but the *full* microsaccade shows a stronger dip than axial alone (Fig. 8B). In the dark-adapted state, the dip depth (as a fraction of the profile peak) is 0 for *none* and *lateral*, 0.036 for *axial* and 0.084 for *full* (Table 4). Resolution therefore depends on axial narrowing and lateral clipping acting together.

**Table 4:** Dip depth of the two-bar response. (fraction of the pooled R1–R6 profile, mean of the UP and DOWN bar sweeps; *r*_*clip*_ = 0.85; dip < 0.02 counts as fused), by pupil state and microsaccade condition. ^*\**^ No reliable dip: the UP bar sweep is fused and the DOWN bar sweep shows a shoulder rather than a dip.

| Pupil state | *none* | *axial* | *lateral* | *full* |
|---|---|---|---|---|
| Dark-adapted | 0 | 0.036 | 0 | 0.084 |
| Halfway | 0 | 0.052 | * | 0.095 |
| Light-adapted | 0.243 | 0.316 | 0.188 | 0.233 |

**Table 5:** Narrowest R1–R6 acceptance angle. Δ*ρ* (^*°*^**) reached during the sweep**, by pupil state and microsaccade condition (*r*_*clip*_ = 0.85; range over R1–R6).

| Pupil state | *none* | *axial* | *lateral* | *full* |
|---|---|---|---|---|
| Dark-adapted | 4.56–5.19 | 4.02–4.75 | 4.22–4.80 | 3.72–4.40 |
| Halfway | 3.69–4.00 | 3.34–3.89 | 3.42–3.70 | 3.09–3.59 |
| Light-adapted | 2.79–2.83 | 2.62–2.80 | 2.58–2.62 | 2.42–2.66 |

##### Pupil adaptation provides the largest effect

Closing the intracellular pupil strips higher-order modes and narrows Δ*ρ* to 55 % to 65 % of its dark-adapted value. Accordingly, the response dip deepens from the dark-adapted to the light-adapted state, and the up/down asymmetry is also clearest in the light-adapted state (Fig. 8B,D; Table 4). Importantly, when light-adapted, the pupil alone is sufficient to resolve the bars: the static (*none*) condition already gives a dip of 0.24, and neither the *full* microsaccade (0.23) nor the *lateral* one (0.19) deepens it, only the *axial* component does (0.32). At the intermediate (*halfway*) state, the full microsaccade opens a dip (0.095) where the static condition is fused. In other words, the spatial benefit of the microsaccade in this model is therefore confined to the dark-adapted and intermediate states, whereas its direction-dependent temporal signature persists in all of them (the lateral offset is 0.16^*°*^ to 0.28^*°*^ across all pupil states). Optical acceptance angles (here 4.56^*°*^ to 5.19^*°*^ at rest for R1–R6 in the forward cartridge) are not directly comparable to voltage-level receptive field (RF) acceptance angles: 6.4^*°*^ to 7.1^*°*^ in the static voltage simulations of [32], and 9.65(106)^*°*^ estimated from their recordings. Voltage-level RFs are wider because of compressive nonlinearities in the photon-to-voltage transformation, which RhabdoForge does not model, and they are also reported to narrow dynamically during microsaccades [31]. The clipping ratio *r*_*clip*_ is a phenomenological factor that makes the ASF narrower during lateral displacement and can thus be read as a stand-in for geometric vignetting, or part of this neural and voltage-level narrowing.

A further caveat concerns what “rest” means here. In the model, rest is the dark-adapted state with the rhabdomere tip at its resting position (17.0 µm), before any pupil mode-stripping, axial narrowing or lateral clipping. Its Δ*ρ* is consistent with the static, dark-adapted optical values of [32] (about 8 % difference when evaluated at the rhabdomere diameters and wavelength they use). It is already close to the value of 4.0^*°*^ (range 3.5^*°*^ to 5.3^*°*^) that [31] take from waveguide theory as the end state of a transient narrowing. RhabdoForge works at the optical level, and thus starts from the optical ASF (i.e. not the voltage-level RF), and the narrowing it produces during a microsaccade is correspondingly small (the full microsaccade narrows R1–R6 to *×* 0.82–0.85 of rest, rather than by nearly half). The phenomenological clipping ratio *r*_*clip*_ is the component that can stand for part of the additional narrowing observed at the voltage level.

Because the readout is a convolution of the stimulus with the ASF, motion cannot recover detail below what the instantaneous ASF resolves: it only converts space into time. Spatial resolution of closely spaced features therefore requires that the ASF actually narrows, and in our model this comes from axial narrowing, aperture clipping, and pupil mode-stripping. By updating the sub-ommatidial geometry on the GPU at each time step, RhabdoForge shows that modelling the eye as an active mechanical filter, rather than a passive camera array, can unlock more accurate neuroethological simulations.

## 4 Discussion

Understanding how insects navigate and interact with the world requires studying their neural circuits in the context of the embodied, dynamic sensory streams that shaped them. In this paper, we introduced RhabdoForge, a modular, hardware-agnostic rendering framework designed specifically to bridge the gap between high-fidelity visual ecology and real-time, closed-loop neuromorphic modelling. By decoupling the sensory morphology from the underlying environment, RhabdoForge allows researchers to construct arbitrary compound eye models, ranging from simple generic lattices to anatomically precise reconstructions of novel species, and place them in any 3D world, from procedural environments to raw LiDAR scans of natural habitats. A central philosophy behind this framework is that the “ommatidium-as-a-pixel” paradigm can be too simplistic for neuroethological modelling. Instead of treating the compound eye as a low-resolution passive camera, RhabdoForge treats it as an active mechanical filter. Sub-ommatidial properties, complex neural superposition wiring, and dynamic photomechanical rhabdomere movements are thus included as fundamental computational elements that pre-process information. The output can then be easily coupled to existing (or new) neuromorphic modelling frameworks and used in agent-based simulation to explore a wide variety of visual behaviours.

This increased realism in representing receptor layout and dynamics still involves some abstraction of the optical processing. For example, RhabdoForge can be contrasted to high-fidelity biophysical models that integrate second order ODEs offline [32, 33] to determine how PIP_2_ cleavage, microvilli refractoriness, stochastic quantum bumps, and Hodgkin-Huxley membranes generate microsaccades and the resulting voltage response. RhabdoForge sits at the algorithmic and input-stream level: given that the microsaccades happen, it asks what light stream actually reaches the agent in a real-time, closed-loop simulation at millisecond temporal resolution, in arbitrarily complex 3D environments. To achieve this on any consumer hardware, RhabdoForge employs an architecture that is loosely inspired by the “matched filters” principle [2, 30]: fast optical reflexes (luminance adaptation, pupil and axial ASF narrowing, and actuation) are computed at the periphery (on the GPU), which thus acts as a computational pre-processor rather than a passive camera, and lower-frequency “central brain” decisions occur on the CPU, in Python. However, the system is designed to be flexible enough to allow, as desired, more mechanistic (rather than phenomenological) modelling of the transformation from light input to neural activation. Either the raw ray-tracing information can be returned from the GPU and processed through any desired model of photoreceptor biophysics, or the dynamic ASF can be computed on the GPU itself, as RhabdoForge already does for the intracellular pupil [81] and, phenomenologically, for lateral aperture-swing clipping [32]. The rendered output thus natively carries a dynamic ASF. Replacing the phenomenological clipping ratio with a mechanistic model of the lens and cone/pigment-cell apertures would be a possible extension, although the computational cost of it might outweigh the small realism gain.

The practical decision to design RhabdoForge as a deterministic radiance model computing purely optical ASFs (e.g. 4.6^*°*^ to 5.2^*°*^ at rest for the *Drosophila* R1–R6 in the forward cartridge), rather than a stochastic model of photoreceptor voltage responses (these are broader: 6.4^*°*^ to 7.1^*°*^ in the voltage simulations of [32], and about 9.7^*°*^ in their recordings), separates optical physics from downstream neural processing. This architectural boundary can directly inform our understanding of biological hyperacuity. Our microsaccade experiment demonstrates that while purely geometric axial displacement can slightly narrow the field and open a shallow spatial dip, the lateral component of the microsaccade effectively smears this spatial profile. Thus, recovering static spatial resolution from the combined microsaccade requires an active narrowing or clipping of the ASF. By explicitly injecting a phenomenological lateral clipping ratio (*r*_*clip*_ = 0.85, calibrated rather than measured) that narrows the ASF during lateral displacement, whether this narrowing arises from geometric aperture interactions (vignetting against the lens and cone/pigment-cell apertures) or from neural and voltage-level nonlinearities downstream of the photon catch, the dark-adapted full microsaccade opens a shallow dip in the two-bar response (0.084 of the profile peak, against 0.036 for the axial component alone). The *r*_*clip*_ ratio was chosen using this same test: it thus demonstrates how the components interact rather than validate the model against true hyperacuity. This allows us to cleanly dissociate the two microsaccade components: the lateral displacement translates the viewing direction to yield direction-dependent temporal coding, while the axial component, coupled with lateral clipping, drives static spatial resolvability. One interesting observation can be made: when light-adapted, the intracellular pupil is already sufficient to resolve the bars, so the spatial benefit provided by microsaccades would only really be useful to the fly when in a dark-adapted state.

The flexibility and utility offered by RhabdoForge are illustrated by the tunnel-centring experiment. First, a complete rendered visual stream can be passed through a minimal correlator array and a proportional controller to generate movement of the agent, closing a complete behavioural loop with latency to spare. Second, because the renderer is downstream agnostic, it is straightforward to substitute alternative processing of the visual stream and test the consequences in identical conditions. In this case, because the Hassenstein-Reichardt correlator reports contrast frequency rather than true angular velocity [95, 96], our first agent inherits a spatial-frequency dependence that real bees do not show: [89] demonstrated that centring is invariant to differences in wall spatial frequency and is displaced only by genuine image motion. When we swapped the HRC with a simple gradient (ratio-based) optic-flow estimator, the agent recovered the bee’s spatial-frequency invariance.

The generality afforded by this modularity includes methods to generate alternative eye models, based on different forms of data for various species. We use the *Drosophila melanogaster* eye to illustrate the full power of these methods, e.g., treating neural superposition as fundamentally a calibration problem. By modelling it as emergent from local geometric rules, we can generate the “imperfect” overcomplete tiling characteristic of fly eyes [97] that is thought to be functional for anti-aliasing. Combined with our stochastic Monte-Carlo sampling and jittering microsaccades, this spatial sampling naturally suppresses spatiotemporal aliasing (Moiré), achieving functionally similar anti-aliasing to more detailed refractory models. The approach frees the model from being a Drosophila-specific tool and opens up design-space exploration. Mixing and matching morphologies, testing evolutionary hypotheses, or grafting a fly’s neural-superposition wiring onto a bee’s high-density lattice, could enhance understanding of how these features match visual tasks across phylogeny.

For example, the ability to produce closed-loop active visual streams invites an investigation into downstream motion processing: RhabdoForge is well positioned to test whether and how physical microsaccade-shaped input reshapes such elementary motion detector tuning, or how the rhabdomere grain’s orientation relative to the global optic flow optimises visual route learning in a navigating agent [98–100]. Similarly, stereopsis can be explored directly, reproducing depth-as-time effects or binocular phase differences for near-versus-far objects would provide a robust secondary validation against active sampling literature [32, 101].

Some limitations should be kept in mind. The lateral clipping ratio *r*_*clip*_ is phenomenological: it was set to produce qualitative effects. Although asset textures and opsin templates can be configured to arbitrary wavelengths, the pipeline currently evaluates light transport across three discrete spectral channels (a constraint inherited from standard RGB display primaries in computer graphics) with a single-wavelength opsin template, rather than simulating a continuous multi-spectral rendering pipeline, and is not a full physical spectral model. Similarly, the current optical model is restricted to apposition-type eyes (including their neural superposition variant), in which each rhabdomere receives light only through its respective ommatidial lens. It thus does not capture optical superposition, as found in the eyes of many nocturnal or crepuscular insects, where a transparent clear zone allows the lenses and cones of many facets to focus light onto a single photoreceptor [21, 61, 102]. Supporting this requires tracing rays through several facets and the clear zone to the receptor layer, and is a target for future work. Finally, with the default sampling settings, the Monte Carlo sampling introduces small errors beyond the expected stochastic noise of a finite ray count. Two are systematic, and arise from how custom ASFs are tabulated: the look-up table has a finite resolution, with a coarser final bin for the outermost *∼*0.4 % of the probability mass, and it only covers the central part of the ASF (by default out to 4*×* its FWHM). This is negligible for the Gaussian and waveguide ASFs, but would bias the signal slightly low for a heavy-tailed custom ASF. The third affects variance: for waveguide ASFs, rays are drawn from a simpler, broader bell-shaped distribution, and each ray is then given a weight that corrects for the difference with the true ASF. Where the true ASF is locally higher than this broader distribution (for example in the tails of higher-order modes), few rays land there but each carries a large weight, so an occasional ray can produce a brief outlier in a rhabdomere’s signal. These weights are not capped by default. Capping them would suppress the outliers at the cost of a small, systematic underestimate of the signal from those tails. At typical ray counts, the first two errors stay well below the Monte Carlo noise.

Beyond addressing these limitations, the system is well positioned to support future expansions on the physics of light, for instance with the addition of a per-rhabdomere e-vector channel for polarisation [103–106], completing a visual ecology suite that already supports spectral sensitivity, colour-bleeding, and soft shadows.

Beyond optics, the architectural decision to expose the Bounding Volume Hierarchy (BVH) to the host CPU opens avenues beyond purely visual research. While RhabdoForge is primarily designed for compound vision, active sensing in insects is fundamentally multimodal, relying heavily on mechanosensation and olfaction [107–118]. By allowing high-speed, CPU-side spatial queries against the same environment used for rendering, the framework provides the necessary infrastructure to simulate tactile modalities via rapid ray- or sphere-casts. This is thus useful for multimodal sensory simulations that aim to replicate the multisensory integration observed in the insect brain.

## 5 Conclusion

RhabdoForge offers a computational framework designed to bridge the gap between high-fidelity visual ecology and real-time neuromorphic modelling. The key advance is to move beyond the traditional assumption of the compound eye as a static array of pixels. Instead the engine models the sensory periphery as an active, mechanical filter. This approach reflects the biological reality that physical actuation, complex structural layouts, and sub-ommatidial dynamics serve as a fundamental pre-processing stage, shaping the visual stream before it reaches the central brain. At the same time, RhabdoForge is built to be a highly accessible tool for the neuroscience and robotics communities, allowing full customisation of the eye design and peripheral processing to the match the biological target and research question, and straightforward integration to existing neural modelling frameworks. By providing a flexible and biologically grounded platform for visual rendering, RhabdoForge serves as a practical foundation for exploring the morphological design space of compound vision, validating hypotheses in visual ecology, and informing bio-inspired sensing strategies for autonomous agents.

## 6 Code availability

The **RhabdoForge** engine and the **PyTinyBVH** library are available as Free and Open-Source Software (FOSS) under MIT license at https://github.com/FlorentLM/rhabdoforge and https://github.com/FlorentLM/pytinybvh.

## 7 Acknowledgements

This work was funded by the UK Research & Innovation Engineering and Physical Sciences Research Council (grant numbers: EP/V008102/1 [recipient BW] and EP/X019632/1 [recipient BW]). The funders had no role in study design, data collection and analysis, decision to publish, or preparation of the manuscript. UK Research & Innovation: https://www.ukri.org/.

For the purpose of open access, the authors have applied a Creative Commons Attribution (CC BY) licence to any Author Accepted Manuscript version arising.

## 9 Supplemental Material

### 9.1 Scene representation and BVH acceleration structure

Like most (if not all) other ray-tracing renderers, RhabdoForge manages the 3D rendering complexity via a multi-level spatial acceleration structure. We use the Bounding Volume Hierarchy (BVH) principle.

While the principles and inner-workings of a BVH are not the scope of this work, it is still useful to mention that we use the state-of-the-art approach of a two-level hierarchy: multiple Bottom-Level Acceleration Structures (BLAS) contain each unique geometric asset in the scene (triangular meshes, point clouds, etc.), and are pre-partitioned into static BVHs, using the Surface Area Heuristic method [119]. A Top-Level Acceleration Structure (TLAS) manages instances of these BLASes. When an instance’s transformation matrix *M* (4 *×* 4) is updated, the TLAS bounding boxes are refitted in *O*(log *N* ) time (avoiding a full tree rebuild), enabling dynamic environments where the insect can interact with moving obstacles or other agents. In other words, the distinction between static BLAS and dynamic TLAS allows the renderer to update the “world” at the same frequency as the eyes’ photomechanical dynamics (see *Sub-ommatidial biophysics and dynamic optics*).

To construct the BVH we use custom Python Bindings (PyTinyBVH [40]) for the lightweight `C++` library TinyBVH [41]. We separate BVH construction and traversal environments: the BVH is built once on the CPU, and the resulting tree data (which includes node bounds and primitive indices) is then uploaded to the GPU in Shader Storage Buffer Objects (SSBOs). The subsequent ray-traversal logic is executed on the GPU within custom GLSL compute shaders.

Unlike traditional triangle-centric engines, the BLAS in our PyTinyBVH bindings and Rhabdo-Forge itself have been designed to treat point clouds as first-class primitives: each point is assigned a radius, allowing the BVH to perform efficient ray-sphere intersections, enabling the rendering of raw environmental datasets such as LiDAR scans or structure-from-motion photogrammetry.

### 9.2 GPU data-structure layout

For efficient memory throughput and cache coherency, RhabdoForge represents eye state in four tightly packed structured arrays on the GPU. These arrays decouple the ommatidium level from the rhabdomere level, and strictly separate static morphological blueprints from dynamic biophysical states, aligned to 16-byte boundaries to satisfy GPU `std430` packing requirements.

At the ommatidium level, the static struct 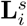 (128 B) encapsulates all fixed lens parameters:

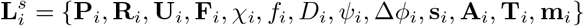

where **P** is the lens position in world space; **R, U, F** form the orthonormal local tangent and optical frame; *χ* is the rhabdomere-bundle yaw; *f* is the lens focal length (the lens-to-rhabdomere lever arm); *D* is the lens aperture; *ψ* is the local hexatic lattice tilt; Δ*φ* are the interommatidial angles; **s** is the local microsaccade actuation axis; **A** holds the lateral and axial microsaccade amplitudes; **T** holds the temporal parameters (ballistic move and return durations, and the fast, slow, pupil and return time constants); and **m** is the local direction of the retinal sheet shift.

The dynamic counterpart **L**^*d*^ (32 B) tracks time-varying states:

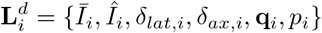

storing the slow (*Ī*) and fast (*Î*) luminance trackers, the lateral (*δ*_*lat*_) and axial (*δ*_*ax*_) displacements, the state of the reflex arc **q** (current phase, time elapsed in it, and fraction of the movement reached at its start), and the pupil closure *p*.

At the rhabdomere level, the static struct 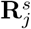 (96 B for the *Drosophila* bundle, which has two bound modes) defines individual rhabdomere parameters:

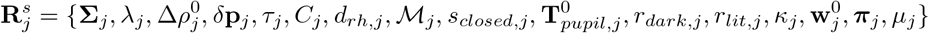

where **Σ** is the 3-channel spectral sensitivity vector; *λ* and *d*_*rh*_ are the peak wavelength and waveguide diameter; 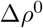 is the resting acceptance angle, stored as a (major, minor) pair 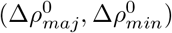 ; *δ***p** is the focal-plane offset; *τ* is the membrane time constant; *C* is the downstream Lamina cartridge index; *M* is a bit-packed metadata identifier (encoding eye index, rhabdomere type, parent ommatidium, and neural-superposition flags); *s*_*closed*_ is the closed-pupil spatial narrowing ratio; 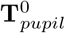 is the 3-channel closed-pupil transmittance vector; *r*_*dark*_, *r*_*lit*_ are the pre-solved axial narrowing ratios under dark- and light-adapted states; *κ∈* [0, 1] is the lateral clipping weight, which grows with the rhabdomere’s offset from the bundle centre (0 on axis); **w**^0^ and ***π*** are arrays of size MAX_LP_MODES storing, for each bound waveguide mode, its resting power weight and its light- to dark-adapted power ratio; and *µ* converts the dimensionless modal half-widths (half the mode’s FWHM) to radians.

Finally, the dynamic rhabdomere struct 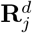 (64 B for two bound modes) maintains the instant-aneous optical state:

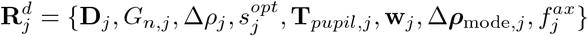

storing the actuated viewing direction **D**, the biochemical gain *G*_*n*_, the dynamic acceptance angle Δ*ρ* (FWHM, also a (major, minor) pair), the axial optical narrowing scale *s*^*opt*^ (which drives photon concentration), the 3-channel dynamic pupil transmittance **T**_*pupil*_, the live mode weights **w** and modal acceptance angles Δ***ρ***_mode_ evaluated by compute shaders at every frame, and the current fraction *f*^*ax*^ of the axial displacement. The size of the mode arrays follows the largest number of bound modes in the rhabdomere bundle.

### 9.3 Random generators

RhabdoForge provides several interchangeable randomness generators to draw sample rays from the ASF-derived PDFs (described in the *Angular sensitivity function* section), balancing stochasticity, hardware performance, and realism.

In the simplest *pseudo-random* mode, the two random variables (*U*_1_, *U*_2_) that condition the ray directions are drawn from a hashed PCG stream [120] seeded per rhabdomere and per sample, yielding fully decorrelated Monte Carlo samples. The remaining modes replace this white-noise sampling with lower-discrepancy alternatives that converge faster for a given *n*, each paired with a per-rhabdomere randomisation so that neighbouring rhabdomeres never share an identical pattern (which would otherwise imprint structured spatial correlations across the array).

Three modes are quasi-random sequences or sets. In *Halton* mode, samples follow the Halton sequence [121] on bases 2 and 3, decorrelated by a per-rhabdomere Cranley-Patterson rotation [122] (a random offset added modulo 1). Because the sequence index advances with each frame, temporal accumulation (see section *Reduction and adaptation*) refines the estimate progressively. The *Hammersley* set [123] is similar to the *Halton* mode, but replaces Halton’s base-2 coordinate with an exactly stratified term 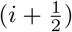, giving lower discrepancy at a fixed, when *n* is known (a marginally cheaper GPU evaluation). It is also decorrelated across rhabdomeres and frames by a Cranley-Patterson shift on both coordinates. Being a fixed-*n* construction it is not extensible, so unlike *Halton* it does not refine progressively across frames. The *Sobol* mode uses a (0, 2)-sequence (van der Corput in base 2 paired with the second Sobol’ dimension [124]), with per-rhabdomere (Owen) scrambling [125, 126] and, as with Halton, a frame-advancing index. It combines the low discrepancy of a (0, 2)-net with progressive refinement, making it the recommended default when the output is temporally integrated. The last two modes are geometric point sets on the disk. In *stratified* mode, the unit square is partitioned into a *g × g* jittered grid (with 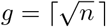 ), each sample is assigned to one cell, and is then randomly perturbed. Because when *n* is not a perfect square the last grid row is only partially populated, the cell assignment is rotated by a per-frame offset, to ensure every direction is covered and the estimator remains unbiased. Finally, the *Fibonacci* mode places samples on a Vogel spiral, which packs points on the disk highly uniformly. A per-rhabdomere, per-frame Cranley-Patterson offset is applied to both the radial and angular coordinates, decorrelating the pattern while preserving its uniformity.

